# Paired activity learning supports systematic identification of mutant-selective EGFR inhibitors

**DOI:** 10.64898/2026.10.21.760684

**Authors:** Jie Zhu, Weikaixin Kong, Liting Huang, Zejia Song, Huiyan Ying, Kari J. Kurppa, Mitro Miihkinen, Su-Jie Zhu, Tero Aittokallio

**Author notes:** Correspondence: Tero Aittokallio, PhD, Institute for Molecular Medicine Finland, FIMM, P.O. Box 20 (Tukholmankatu 8), FI-00014 University of Helsinki, Helsinki, Finland. Competing interests: MM and TA have received unrelated research funding from Mobius Biotechnology GmbH. The other authors declare that they have no competing interests.

## Abstract

Acquired resistance to epidermal growth factor receptor (EGFR) inhibitors remains a major clinical challenge in non-small cell lung cancer (NSCLC), where successive rounds of therapy with first- and third-generation EGFR inhibitors often lead to increasingly resistant EGFR variants. In particular, EGFR L858R/T790M/C797S triple mutation renders tumors resistant to earlier-generation inhibitors, while disrupting the covalent engagement required by third-generation inhibitors such as osimertinib. Here, we developed an integrated computational–experimental framework for identifying compounds with differential activity between the triple-mutant EGFR and wild-type EGFR. Matched bioactivity measurements were leveraged to train a mutant-aware paired-activity regression framework, and the predicted candidates meeting stringent potency and mutant-preference criteria were subsequently evaluated using a support vector machine classifier for triple-mutant activity. Application of this sequential screening approach to 12 million molecules identified 16 consensus hits, including several previously reported inhibitors of drug-resistant EGFR variants. Based on their predicted differential activity and degree of prior characterization, five hit compounds were prioritized for initial biochemical evaluation, and two were selected for further biochemical, cellular and structural characterization. Mutant EGFR crystal structures in complex with the best-performing compounds revealed two distinct noncovalent binding modes within the ATP-binding cleft, demonstrating the ability of the framework to identify potent inhibitors with diverse binding modes. Together, this work illustrates how incorporating mutant-selectivity into a predictive framework as paired activities support the identification of mutant EGFR inhibitors and provide insights into their potency and selectivity. The discovered complementary structures provide experimentally defined templates for optimizing potency, mutant preference and drug-like properties of these and future compounds combating drug resistant NSCLC.

**Significance:** This work introduces a generally-applicable computational–experimental learning framework for systematic screening of mutant-selective inhibitors and provides a proof-of-concept for selective targeting of resistance-driving mutations using paired-activity learning and predictive modelling.

## Introduction

Non-small cell lung cancer (NSCLC) accounts for approximately 85% of lung cancers^1^ and includes a clinically important subgroup defined by EGFR mutations^2^. Activating mutations such as exon 19 deletions and L858R confer sensitivity to EGFR tyrosine kinase inhibitors (TKIs)^3–6^, but this gained activity is frequently compromised by acquired resistance. An early genetic resistance-mechanism is the emergence of T790M mutation, either acquired or selected during treatment with earlier-generation inhibitors, which increases ATP affinity and reduces TKI effectiveness^7,8^. The third-generation TKI osimertinib^9^ was developed to overcome this resistance through covalent binding to cysteine 797 (C797) within the EGFR ATP-binding pocket. Despite its clinical efficacy, resistance to osimertinib frequently emerges, with substitution of this critical cysteine by serine (C797S) representing an important on-target mechanism that prevents covalent drug binding^10^. Importantly, C797S mutation is the dominant resistance mechanism to osimertinib, highlighting the need for next-generation TKIs overcoming the resistance caused by the triple mutation profile (L858R/T790M/C797S)^10,11^.

Current therapeutic approaches to C797S-containing EGFR include allosteric inhibition, reversible ATP-site inhibition and drug combinations that engage distinct sites^12–16^. For instance, the allosteric inhibitor EAI045 required cetuximab combination treatment for boosting its tumor suppression efficacy in the reported preclinical models^12^. Across the strategies attempting to target mutant EGFR, a central challenge is the lack of systematic approaches capable of predicting molecular affinities that favor resistant EGFR mutants over wild-type (WT) EGFR. As WT EGFR inhibition contributes to dose-limiting adverse effects, including rash and diarrhea, and consequently narrows the therapeutic window^17^, such approaches could provide early insights into mutant-specific potency and selectivity, enabling more informed lead candidate prioritization^18,19^. Although virtual screening can prioritize lead compounds, predicting mutant activity alone does not establish a mutant preference. We therefore hypothesized that an integrated paired mutant and WT EGFR activity profiling and modelling across EGFR variants is needed to jointly predict both mutant-selective potency and differential activity.

Here, we combined a mutant-aware paired-activity regression framework that predicts EGFR mutant activity, WT activity and their activity difference, with a separately trained EGFR L858R/T790M/C797S support vector machine (SVM) activity classifier. Candidate compounds were required to satisfy pre-defined potency criteria and exhibit concordant mutant preference based on the difference between predicted mutant and WT activities. Screening more than 12 million molecular records, followed by candidate auditing and prioritization, yielded five compounds for experimental evaluation. Biochemical and cellular profiling identified two structurally distinct inhibitors with a low-nanomolar affinity for triple-mutant EGFR, one demonstrating approximately 1,000-fold preferential binding to mutant over WT EGFR. Structural analyses using X-ray crystallography further revealed distinct noncovalent binding modes, providing a foundation for subsequent lead optimization against drug-resistant NSCLC. Together, this work provides a proof-of-concept for selective targeting of cancer-driving mutations using paired-activity learning and predictive modelling.

## Results

### Integrated dataset of wild type and mutant EGFR activities for paired activity modelling

To establish paired activity predictors of mutant EGFR inhibition with a reduced affinity towards wild-type receptor, we first integrated and curated a paired-activity dataset consisting of 1,223 matched WT–mutant measurements, representing 810 unique compounds and 30 EGFR variant settings (**Table 1**; **Supplementary Table 1**; **Supplementary Figure 1a–c**). Most activity records were obtained exclusively from ChEMBL release 37 (1,189 records), with smaller contributions from BindingDB (28) or both databases (6). The harmonized dataset comprised 1,094 IC_50_, 81 K_d_, and 48 K_i_ measurements. More detailed distributions of data sources, activity endpoints, assay classes, confidence levels and variant types are provided in **Supplementary Table 2**.

**Table 1.** Composition and use of the paired-regression and triple-mutant classification datasets.

| Usage | Paired regression dataset | Triple-mutant classification dataset |
| --- | --- | --- |
| Prediction task | Continuous WT, mutant and selectivity regression | Triple-mutant active/inactive classification |
| Modeling records | 1,223 | 2,099 |
| Unique compounds | 810 | 2,099 |
| EGFR variant settings | 30 | L858R/T790M/C797S only |
| Triple-mutant paired records | 84 | Not applicable |
| WT endpoint | Continuous pActivity(WT) | Not included |
| Mutant endpoint | Continuous pActivity(mutant) | Binary activity label |
| Selectivity endpoint | $\Delta pActivity = \text{mutant} - \text{WT}$ | Not directly modeled |
| Activity types | $IC_{50}$, $K_d$ and $K_i$ | $IC_{50}$ |
| Active definition | Continuous measurement | Measured $IC_{50} < 50$ nM |
| Inactive definition | Continuous measurement | 339 measured $IC_{50} > 500$ nM + 763 presumed negatives |
| Data sources | ChEMBL37 and BindingDB | ChEMBL, BindingDB and patent literature |
| Primary validation | Repeated scaffold-based 5-fold CV | Repeated scaffold-based 5-fold CV |

Data are reported after curation and compound standardization. $pActivity = -\log_{10}(\text{activity in mol l}^{-1})$ . $\Delta pActivity = \text{mutant } pActivity - \text{WT } pActivity$ . The classifier negative class comprised 339 compounds with measured $IC_{50} > 500$ nM and 763 presumed-negative compounds selected from ChEMBL.

EGFR double-mutant T790M/L858R was the most frequently represented variant setting, with 564 paired WT-mutant activity measurements (46.12%), whereas only 84 paired measurements (6.87%) were available for the EGFR triple-mutant L858R/T790M/C797S (**Supplementary Table 1**). The measured WT and mutant activities covered broad and partly overlapping ranges, with subsets of compounds displaying either mutant-preferential (52.82%) or WT-preferential (9.65%) activity (**Supplementary Figure 1d**).

Due to the low number of triple-mutant records, we leveraged the paired measurements across 30 EGFR variants in regression model development (**Supplementary Table1**). We separately assembled a triple-mutant classification dataset from ChEMBL, BindingDB and patent literature. The dataset contained binary activity labels for 2,099 unique compounds, comprising 997 measured actives with IC_50_ values below 50 nM, 339 measured inactives with IC_50_ values above 500 nM and 763 compounds randomly sampled from ChEMBL as presumed-negative examples (**Table 1**; **Supplementary Figure 1e–f**). Because the proportions of active and inactive compounds differed among ChEMBL, BindingDB and patent-derived records, we evaluated data source-controlled classifier performance (**Supplementary Table 3**).

### Protein language model representations improve paired EGFR activity prediction

Next, we developed and systematically evaluated a predictive framework to prioritize small-molecule compounds with potent activity against EGFR L858R/T790M/C797S and reduced activity against WT EGFR (**Figure 1**). We compared various molecular fingerprints, mutation representations and machine-learning approaches, including support vector regression (SVR), multitask multilayer perceptrons (MLP) and graph isomorphism networks (GIN), across three prediction tasks: mutant pActivity, WT pActivity and their difference (ΔpActivity). Based on these comparisons, we established a two-stage prioritization workflow combining an ensemble preference with a separately trained triple-mutant activity support vector machine (SVM) classifier as a secondary filter.

**Figure 1:**
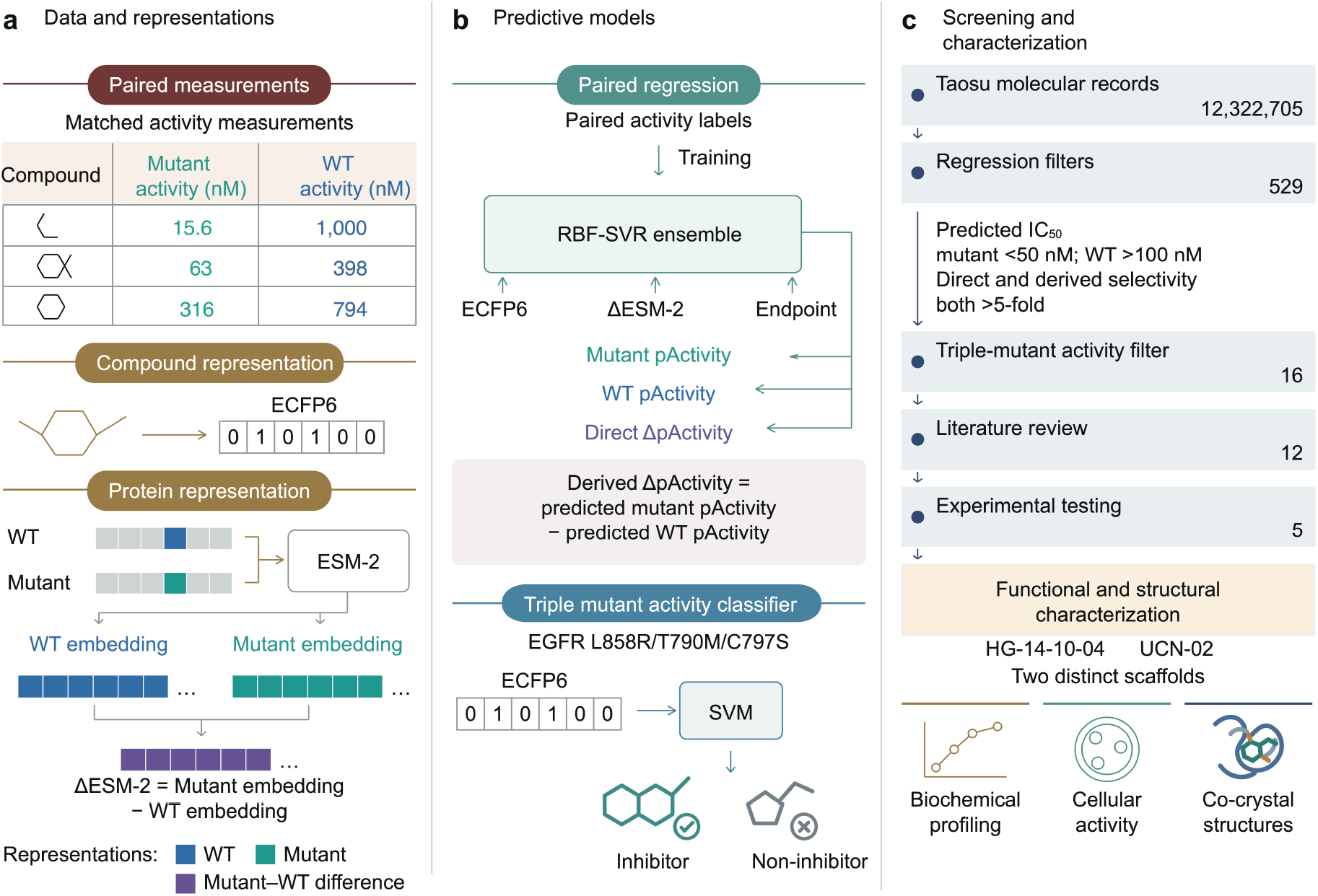
A mutant-aware paired activity regression–classification cascade prioritizes compounds with predicted mutant preference. **a**, Compound and protein representations used for paired EGFR activity modelling. Compounds were encoded as Morgan fingerprints (ECFP6) and linked to matched wild-type (WT)-mutant activity measurements. EGFR variants were represented using mutation-aware protein features derived from the difference between the variant and WT ESM-2 protein language model^20^ embeddings (ΔESM-2). The paired endpoints were mutant pActivity and WT pActivity, modelled using separately trained regressors. **b,** Paired activity prediction of potency and mutant preference for candidate screening. Five RBF-SVR models were trained for the three endpoints (mutant pActivity, WT pActivity and direct ΔpActivity) using the training partitions of a frozen scaffold-based five-fold split, and their predictions were averaged within each endpoint. A separate, derived ΔpActivity estimate was calculated by subtracting predicted WT pActivity from predicted mutant pActivity. Compounds were retained when the predicted mutant IC_50_ <50 nM, the predicted WT IC_50_ >100 nM, and both direct and derived estimates indicated >5-fold mutant preference. The retained compounds were subsequently evaluated using the ECFP6-SVM triple mutant-activity classifier. **c,** Screening funnel from the Taosu virtual library to experimental evaluation. Of 12,322,705 molecular records, 529 standardized, non-duplicate structures satisfied the regression criteria, of which 16 were further classified as triple-mutant active. Four compounds overlapping with the model-development datasets were not advanced to experimental evaluation. Among the remaining 12 compounds, four with previously reported biochemical and cellular activity in C797S-containing or triple-mutant EGFR models were retained as literature-supported positive hits but not prioritized for additional testing. Three of the remaining eight candidates were not advanced because of structural redundancy or commercial unavailability, yielding five compounds for initial biochemical evaluation. HG-14-10-04 and UCN-02 were subsequently selected for detailed biochemical, cellular and structural characterization.

First, we systematically evaluated how various compound, mutation and model representations affected the prediction of mutant activity and EGFR state selectivity. All configurations were evaluated on the same 1,223 paired measurement records, representing 810 unique compounds, using identical cross-validation with scaffold-based five-fold partitions across the same ten random seeds (**Figure 2a**). Compounds sharing a Bemis–Murcko scaffold were assigned to the same fold, preventing scaffold overlap between the training and held-out sets, which led to a relatively challenging prediction task, compared to random cross-validation (CV).

**Figure 2:**
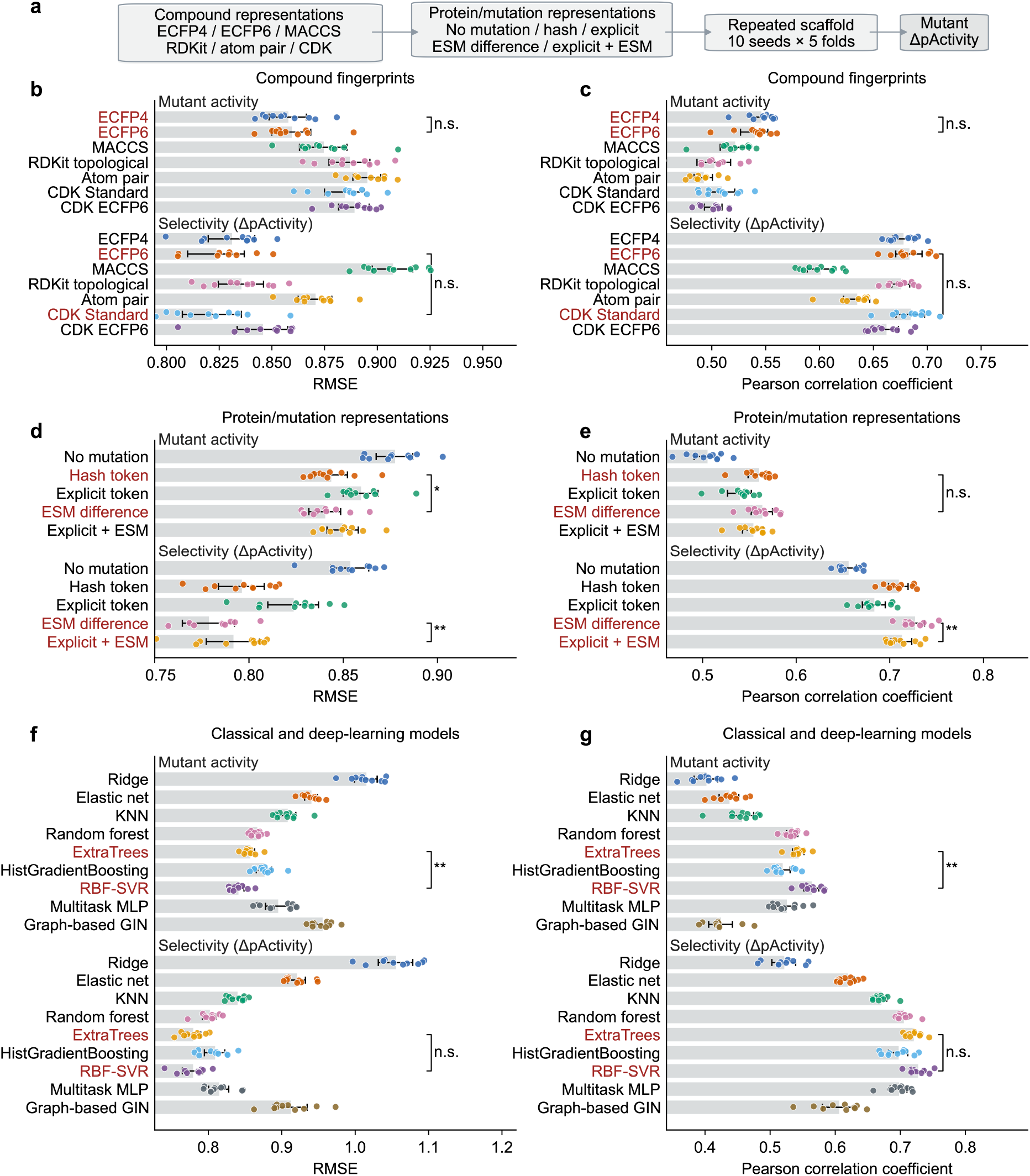
Mutant-aware representations improve EGFR mutant activity and selectivity predictions of paired activity models. **a**, Benchmarking workflow. A total of 810 compounds were encoded using seven two-dimensional molecular representations and combined with alternative EGFR variant representations. All models were evaluated using identical repeated scaffold-based five-fold cross-validation across ten random seeds. Primary endpoints were mutant pActivity and selectivity (ΔpActivity). **b**,**c**, Comparison of molecular fingerprints using support vector regression (SVR) with the explicit mutation descriptor. **d**,**e,** Comparison of mutation representations with ECFP6 as the fixed compound representation. **f**,**g**, Comparison of seven classical regression algorithms and two deep-learning architectures including a multitask multilayer perceptron (MLP) and a graph isomorphism network (GIN). The bars represent the average of the seed-level five-fold averages across ten repeated scaffold partitions, error bars are 95% confidence intervals and points correspond to individual seed-level averages. The red labels indicate the two highest-performing configurations for the corresponding endpoint and evaluation metric. Note that the scale of x-axis varies between the plots. Statistical comparisons were based on two-sided Wilcoxon signed-rank tests across the same ten seeds with the Holm correction. *p < 0.05, **p < 0.01, ***p < 0.001; n.s., not significant. RMSE, root mean square error.

We benchmarked seven molecular fingerprints covering circular, substructure-key, topological and atom-pair representations^21–24^, which allowed us to assess how different encodings of compound structure affected activity and selectivity prediction (**Figure 2b**, **c**). ECFP4 and ECFP6 led to similar root mean square error (RMSE) values and Pearson correlations. ECFP6 outperformed CDK Standard fingerprints for mutant and WT activity prediction, while their direct ΔpActivity performance was nearly identical (**Supplementary Figure 2**). We therefore selected ECFP6 as a common molecular representation for the regression modelling, because it performed consistently well across the activity prediction and the three selectivity tasks.

To represent different EGFR mutational states (**Supplementary Table 1**), we calculated the difference between variant and WT sequence embeddings generated by the ESM-2 protein language model (ΔESM-2)^20^. This representation encodes mutation-associated differences in sequence context, allowing the regression models to consider EGFR variant information alongside compound structure. Incorporating ΔESM-2 improved prediction relative to the explicit mutation descriptor, with the largest gains observed for ΔpActivity prediction (**Figure 2d**, **e)**. Relative to the explicit mutation descriptor, ΔESM-2 reduced mutant-activity RMSE and increased Pearson correlation, with the largest improvement for ΔpActivity prediction. Combining explicit mutation descriptors with ΔESM-2 did not further improve prediction performance (**Figure 2d**, **e**).

To assess how different learning architectures perform in the paired mutant-WT activity prediction tasks, we compared nine models spanning linear regression, nearest-neighbour methods, tree ensembles, kernel methods^25^, together with a multitask multilayer perceptron^26^ and a graph isomorphism network (GIN)^27^ (**Figure 2f**, **g**). This comparison examined whether multitask neural networks and molecular graph learning offered advantages over conventional regression algorithms within the paired mutant–WT prediction framework. Although multiple models performed well, especially for the ΔpActivity prediction, ExtraTrees (Extremely randomized trees) and RBF-SVR (Radial Basis Function Support Vector Regression) showed the best performance. We therefore selected ECFP6–ΔESM-2 support vector regression (SVR) for mutant activity and selectivity prediction, and ECFP6-based regression for WT activity prediction.

Secondary metrics, including mean absolute error and Spearman correlation, supported the same overall ranking (**Supplementary Figure 3**). For each paired measurement record, we also averaged its out-of-fold (OOF) predictions across the ten scaffold partitions and recalculated performance (**Supplementary Figure 4**). These repeated-CV-averaged OOF summaries differed from the average seed-level metrics, because they were calculated after averaging predictions, and therefore represent record-level consensus performance across the repeated cross-validation procedure, rather than performance from an individual scaffold partition. These results summarize prediction–measurement agreement across the curated paired dataset.

### Scaffold and data source controls improve triple mutant activity classification

Since the EGFR triple-mutant measurements represented only a small fraction of the paired-regression dataset, we complemented the continuous activity and selectivity predictions from the RBF-SVR models with a separately trained triple-mutant activity classifier. This classifier used a larger dataset of EGFR L858R/T790M/C797S activity labels to distinguish active from inactive compounds. We therefore benchmarked molecular fingerprints and classification algorithms for this binary prediction task (**Figure 3a**). Among seven fingerprints evaluated with a common SVM classifier and matched scaffold partitions, ECFP6 achieved the highest ROC–AUC and Matthews correlation coefficient (MCC), with significant differences compared to the next-ranked fingerprint across matched seeds (Figure 3b-c).

**Figure 3:**
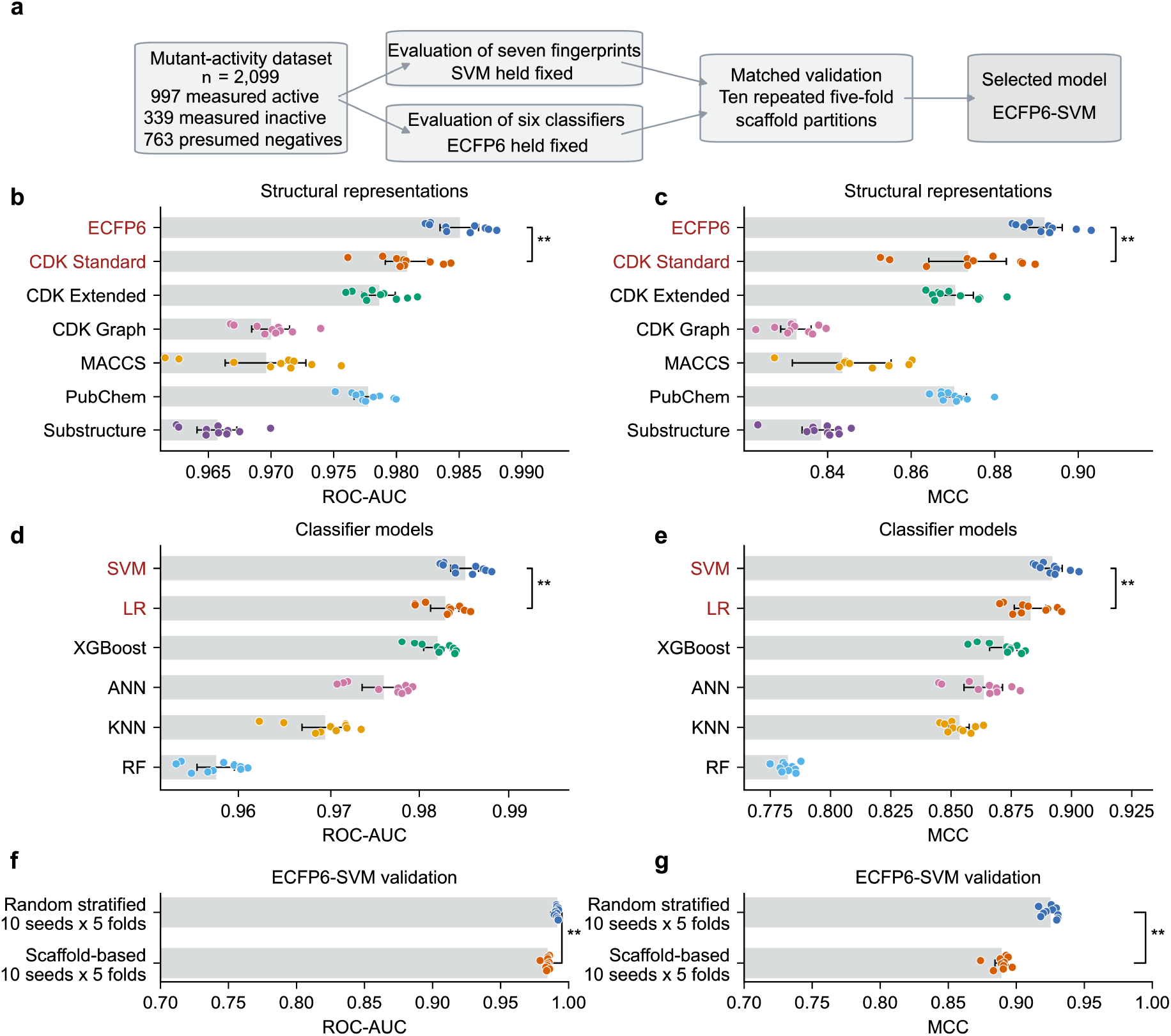
Repeated scaffold validation identifies ECFP6-SVM best for triple-mutant activity classification. **a**, Benchmarking of molecular-fingerprints and classifiers with repeated scaffold validation for the EGFR L858R/T790M/C797S activity classification. **b**,**c**, Performance of SVM classifiers trained with structural fingerprints. **d**,**e**, Performance of SVM, logistic regression (LR), XGBoost, artificial neural network (ANN), k-nearest neighbours (KNN) and random forest (RF) models using ECFP6 as the fixed molecular representation. **f**,**g**, Comparison of repeated random-stratified and scaffold-based five-fold cross-validation for the selected ECFP6-SVM model, evaluated by ROC-AUC and MCC. The bars represent means, error bars are 95% confidence intervals and points represent individual seed-level five-fold averages. The red labels indicate the two highest-performing configurations for the corresponding endpoint and evaluation metric. Note that the scale of x-axis varies between the plots. Fingerprint and algorithm comparisons were based on two-sided Wilcoxon signed-rank tests across ten matched scaffold-split seeds with the Holm correction. *p < 0.05, **p < 0.01, ***p < 0.001. ROC-AUC, area under the receiver operating characteristic curve. MCC, Matthews correlation coefficient.

With ECFP6 fingerprints fixed, SVM ranked best among the six tested classification algorithms both in terms of ROC-AUC and MCC (**Figure 3d**, **e**). Logistic regression and XGBoost followed closely behind, while the artificial neural network, k-nearest neighbors and random forest ranked lower. The high performance of SVM was supported by secondary metrics, including balanced accuracy, precision and recall (**Supplementary Figure 5**). We therefore selected ECFP6–SVM as the triple-mutant activity classifier based on its overall performance across the evaluated metrics.

We further validated the selected triple mutant activity classifier under two training-test split settings. As expected, the randomly stratified cross-validation resulted in higher accuracy, compared to scaffold-based validation (**Figure 3f,g**). The lower scaffold-split performance is consistent with a more demanding prediction setup, in which compounds sharing a Bemis–Murcko scaffold are kept in the same CV fold. Performance nevertheless remained high under this validation setup as well.

To assess whether classification performance was influenced by differences in class composition between data sources, we evaluated the ECFP6-SVM triple-mutant activity classifier across individual sources and after balancing active and inactive compounds within each source (**Supplementary Figure 6**). The classifier retained its strong performance within the ChEMBL and patent-derived subsets and after balancing the classes within each source. These controls over data source and molecular scaffolding support its use as a complementary triple-mutant activity filter within the screening framework.

### Paired-activity regression-classification cascade prioritizes mutant-active compounds

The optimal activity screening cascade leveraged five-model SVR ensembles for mutant activity, WT activity and direct ΔpActivity prediction (**Figure 1b**, **c**). Successful compound candidates had to satisfy four regression criteria: predicted mutant IC_50_ <50 nM, predicted WT IC_50_ >100 nM, directly predicted mutant preference >5-fold and derived mutant preference ΔpActivity >5-fold. Agreement between the two selectivity estimates served as an internal consistency requirement; because the regressors used the same data, however, the estimates were not treated as independent validation.

Screening the Taosu chemical collection (>12M virtual compounds)^28^ yielded 529 standardized, non-duplicate structures that satisfied the regression-based potency and mutant-preference criteria (**Figure 1c**). These compounds were further assessed using ECFP6–SVM triple-mutant activity classifier, which classified 16 as L858R/T790M/C797S active (**Supplementary Table 5**). The resulting consensus candidates therefore combined regression-predicted potency and mutant preference with a triple-mutant activity classification, and were advanced to candidate auditing and prioritization.

The 16 consensus hits were reviewed for overlap with the model-development datasets and previously reported EGFR activity. Four compounds had exact molecular-connectivity matches to the model-development datasets and were not advanced for experimental evaluation. Among the remaining 12 hits, TQB3804^11^, OBX02-011^29^, CZC-54252^30^ and NVP-TAE684^31^ had published biochemical and cellular activity in C797S-containing or triple-mutant EGFR models. Their recovery supported agreement between the screening predictions and existing experimental evidence, although these compounds were not prioritized for additional testing.

Further prioritization based on characterization gaps, structural redundancy and experimental tractability yielded five compounds for initial biochemical evaluation: (S)-sunvozertinib, UCN-02, HG-14-10-04, GSK1904529A and Mps1-IN-2 (**Supplementary Figure 7**). HG-14-10-04 had previously been examined as an intermediate in the medicinal-chemistry series reported by Chen et al.^16^, who demonstrated its biochemical activity against EGFR L858R/T790M/C797S. However, we identified no prior report of its cellular activity in the exact triple-mutant setting or its EGFR-bound crystal structure. These characterization gaps motivated their inclusion for further investigation.

### Biochemical and cellular profiling reveals distinct EGFR activity profiles for HG-14-10-04 and UCN-02

To establish biochemical profiling conditions, we first characterized the ATP dependence of recombinant EGFR L858R/T790M/C797S over an ATP concentration range of 0–128 µM. Michaelis–Menten analysis yielded a K_m_ of 6.7 µM (**Supplementary Figure 8a**). We next performed an initial candidate-profiling experiment in which all five selected compounds were tested across a broad concentration range at 500 µM ATP. Under these conditions, (S)-sunvozertinib, HG-14-10-04 and UCN-02 produced concentration-dependent inhibition, whereas GSK1904529A and Mps1-IN-2 showed little or no inhibition over the tested concentration range (**Supplementary Figure 8b**). HG-14-10-04 and UCN-02 showed strongest inhibition among the candidates, and were therefore selected for further biochemical, cellular and structural characterization (**Figure 4a**). In confirmatory assays conducted at 1,000 µM ATP, well above the measured ATP K_m_ of 6.7 µM, HG-14-10-04 and UCN-02 inhibited EGFR L858R/T790M/C797S with IC_50_ of 3.1 µM and 5.9 µM, respectively (**Figure 4b**). Assuming ATP-competitive inhibition, application of the Cheng–Prusoff relationship yielded estimated K_i_ values of approximately 20.6 and 39.3 nM, respectively, supporting nanomolar inhibitory potency.

**Figure 4:**
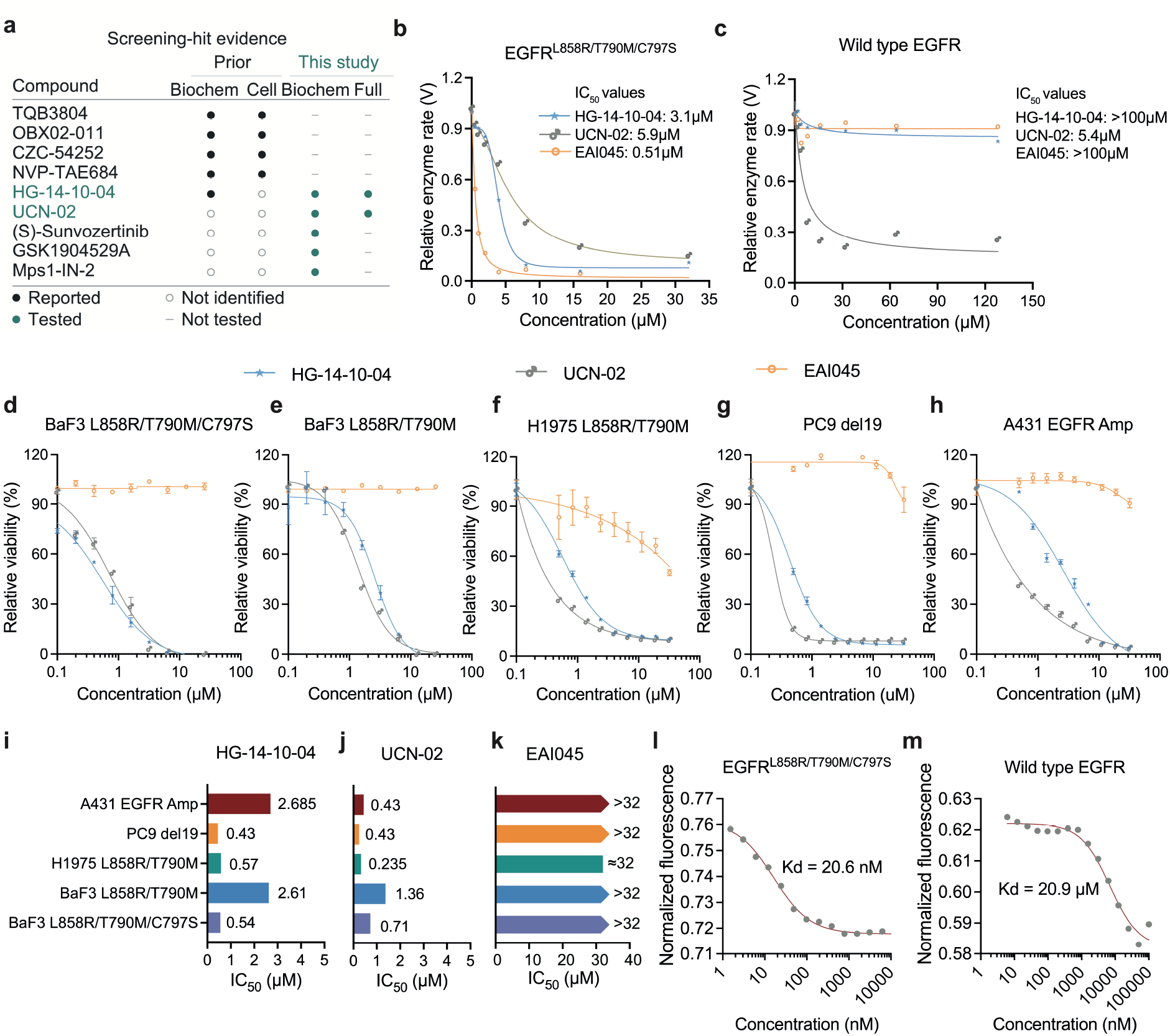
Biochemical and cellular profiling identifies HG-14-10-04 and UCN-02 as active EGFR-mutant inhibitors. **a**, Prior experimental evidence and follow-up testing for selected regression–classification consensus hits. Filled black circles indicate reported biochemical or cellular activity in relevant C797S-containing or triple-mutant EGFR models; open circles indicate that no corresponding evidence was identified in the literature review, not demonstrated inactivity. Green colored circles indicate initial biochemical testing or full biochemical, cellular and structural profiling performed in this study. Complete report of the 16 consensus hits is provided in Supplementary Figure 7. **b**,**c**, Biochemical dose–response profiles of HG-14-10-04,UCN-02 and EAI045 against recombinant EGFR L858R/T790M/C797S and wild-type (WT) EGFR at 1,000 µM ATP. EAI045 was included as a reference, selective inhibitor of the L858R/T790M mutant. **d**–**h**, Cellular dose–response profiles of HG-14-10-04, UCN-02 and EAI045 in Ba/F3 cells expressing EGFR L858R/T790M/C797S (**d**), Ba/F3 cells expressing EGFR L858R/T790M (**e**), H1975 cells harboring endogenous EGFR L858R/T790M (**f**), PC9 cells carrying an EGFR exon 19 deletion (**g**) and A431 cells with amplified WT EGFR (**h**). **i**-k, Summary of the fitted cellular IC_50_ values across the five cellular models. Dose-response data are presented as mean ± SEM from three replicate measurements per concentration. Curves were fitted with nonlinear regression. **l,m,** Microscale thermophoresis (MST) measurements of HG-14-10-04 binding to EGFR L858R/T790M/C797S and WT EGFR, with apparent K_d_ values of 20.6 nM and 20.9 µM, respectively.

Interestingly, HG-14-10-04 and UCN-02 exhibited clearly different activity profiles against WT EGFR. HG-14-10-04 caused little WT EGFR inhibition over the tested concentration range, similar to EAI045, a known selective inhibitor of the L858R/T790M mutant, whereas UCN-02 substantially reduced WT EGFR activity (**Figure 4c**). Thus, the enzyme assays supported a mutant-preferential profile for HG-14-10-04, whereas the triple-mutant activity of UCN-02 was accompanied by considerable WT EGFR inhibition.

We next compared the cellular activities of HG-14-10-04 and UCN-02 across a panel of EGFR-dependent cell models (**Figure 4d–h**). Even though there were differences in IC_50_ levels, both compounds inhibited the viability of Ba/F3 cells expressing EGFR L858R/T790M/C797S (**Figure 4d**), Ba/F3 cells expressing EGFR L858R/T790M, (**Figure 4e**), H1975 cells harbouring endogenous EGFR L858R/T790M (**Figure 4f**), and PC9 cells carrying an EGFR exon 19 deletion (**Figure 4g**), much stronger than the reference inhibitor EAI045 (**Figure 4i-k**). These findings show that neither compound was exclusively active against the triple mutant only and that both retained activity across EGFR-mutant cell models with distinct, literature-reported sensitivity and resistance profiles to established and investigational EGFR inhibitors (**Supplementary Table 6**).

A clearer difference between the two compounds emerged in A431 cells, which harbor amplified wild-type EGFR. HG-14-10-04 inhibited A431 cell viability with an IC_50_ of 2.685 µM, approximately fivefold higher than its IC_50_ in triple-mutant Ba/F3 cells. By contrast, UCN-02 inhibited A431 cells with an IC_50_ of 0.43 µM, comparable to or lower than its activity in the tested EGFR-mutant cell models (**Figure 4h–j**). These cellular results are consistent with a mutant-preferential activity profile for HG-14-10-04, whereas UCN-02 showed activity across both EGFR-mutant and WT-EGFR-amplified cell models. Finally, microscale thermophoresis measurements indicated substantially stronger apparent binding of HG-14-10-04 to EGFR L858R/T790M/C797S than to wild-type EGFR, with K_d_ values of 20.6 nM and 20.9 µM, respectively, indicating a 1000-fold selectivity (**Figure 4l, m**). Together, the biochemical, cellular and binding measurements support a mutant-preferential profile for HG-14-10-04.

The experimental assays supported triple-mutant activity of both compounds with a clearer biochemical mutant preference for HG-14-10-04. The WT activity of UCN-02 showed that passing the computationally-predicted selectivity criteria does not necessarily ensure experimental WT sparing. We next determined how these two compounds engaged the resistant EGFR ATP-binding pocket.

### Structural analyses reveal distinct noncovalent binding modes

To determine how structurally distinct compounds engage drug-resistant EGFR, we determined crystal structures of the EGFR T790M/C797S kinase domain in complex with HG-14-10-04 and UCN-02 (**Figure 5**; **Table2**). Co-crystallization was initially attempted using the EGFR L858R/T790M/C797S kinase domain employed in the biochemical assays; however, the resulting diffraction data were of insufficient resolution for reliable analysis of ligand–protein interactions. Comparison with these low-resolution data indicated that the overall kinase fold and the principal features of the ATP-binding pocket were conserved in the T790M/C797S construct, which reproducibly yielded higher-resolution diffraction data. We therefore used the T790M/C797S construct to define the ligand-binding modes. This strategy is consistent with several previous studies^12,32^ in which inhibitor activity was evaluated against L858R/T790M/C797S EGFR, whereas T790M/C797S or related crystallization-compatible constructs were used for structural analysis. In both structures, the ligands occupied the ATP-binding cleft between the N- and C-terminal lobes without inducing a major rearrangement of the overall kinase fold.

**Figure 5:**
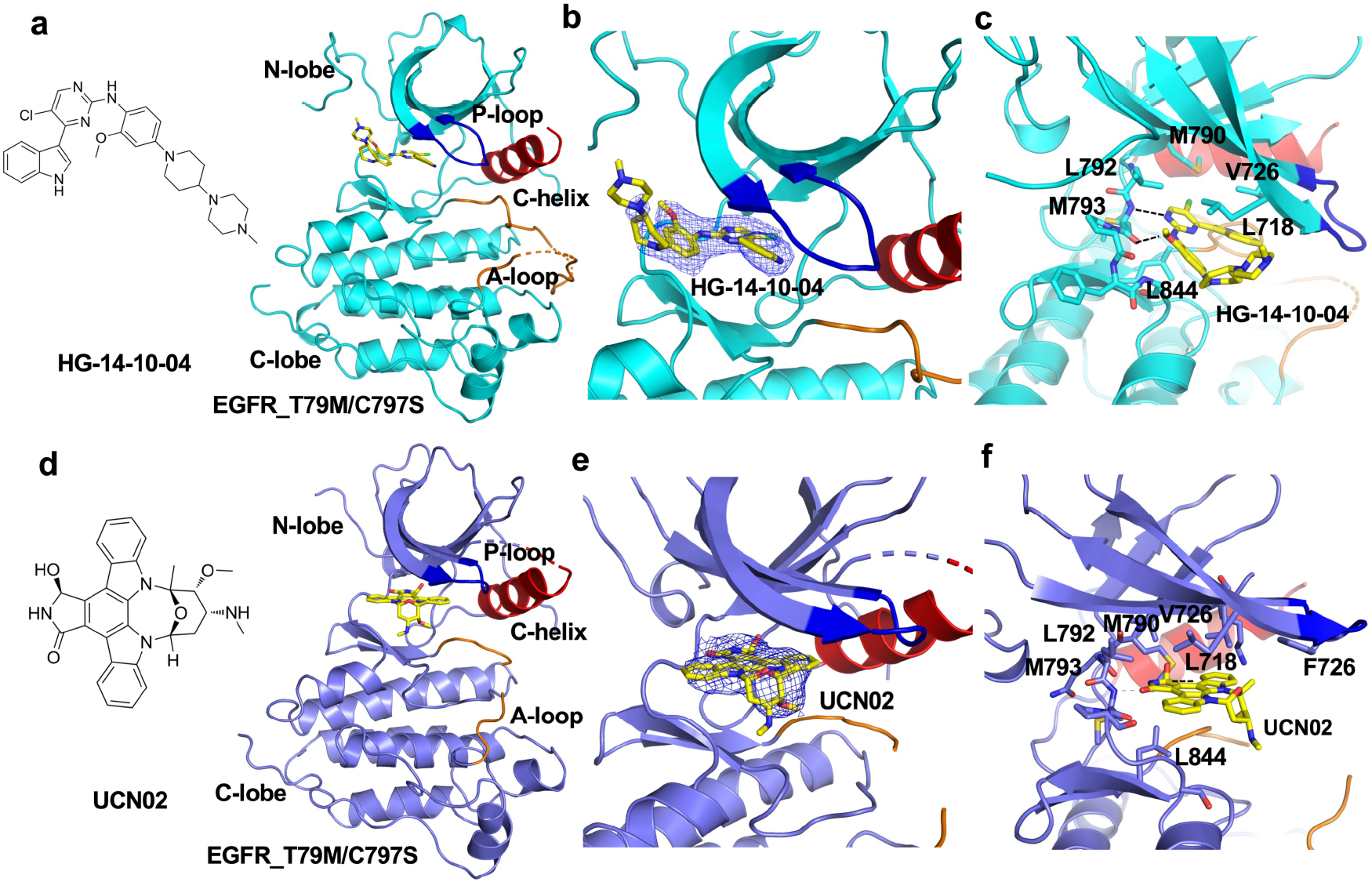
HG-14-10-04 and UCN-02 adopt distinct noncovalent binding modes in EGFR T790M/C797S. **a-c**, Crystal structure of the EGFR T790M/C797S kinase domain in complex with HG-14-10-04 (PDB ID: 46VO). **a**, Chemical structure of HG-14-10-04 and overall view of the kinase-ligand complex. The N-lobe, C-lobe, P-loop, alphaC-helix and activation loop (A-loop) are indicated. **b**, Close-up view of HG-14-10-04 in the ATP-binding cleft with the ligand omit map shown as a blue mesh (2Fo-Fc, contoured at 1.0 sigma). **c,** Ligand-protein contacts involving residues surrounding HG-14-10-04; dashed lines indicate hydrogen bonds. **d-f**, Crystal structure of EGFR T790M/C797S in complex with UCN-02 (PDB ID: 46VR). **d**, Chemical structure of UCN-02 and overall view of the complex. **e**, Close-up view of UCN-02 in the ATP-binding cleft with the corresponding ligand omit map shown as a blue mesh (2Fo-Fc, contoured at 1.0 sigma). **f**, Residues surrounding UCN-02 and the associated polar and hydrophobic contacts; dashed lines indicate hydrogen bonds. Ligands are shown as yellow sticks.

HG-14-10-04 occupied the ATP-binding pocket in an extended conformation (**Figure 5b, c**). Its heteroaromatic core inserted into the adenine-binding region, whereas its basic terminal substituent extended toward the solvent-exposed entrance of the pocket. The central scaffold was supported by well-defined density in the 2Fₒ−Fc omit map, while a weaker density around the terminal group suggested conformational flexibility in this region. HG-14-10-04 formed two hydrogen bonds with the hinge residue Met793. Its aromatic scaffold also contacted Leu718, Val726, Met790, and Leu844 through hydrophobic interactions. The contact between its chlorine substituent and Met790 provides a plausible structural hypothesis for recognition of the T790M-containing pocket, although the contribution of this interaction to mutant preference remains to be established experimentally.

UCN-02 occupied the same ATP-binding cleft but adopted a distinct orientation consistent with its more rigid polycyclic scaffold (Figure 5d-f). Ligand heteroatoms formed hinge-region hydrogen bonds with Met793, while hydrophobic contacts with Leu718, Phe723, Val726, Met790, Leu792, Met793, and Leu844 stabilized the complex. Continuous density in the corresponding omit map supported the modeled UCN-02 binding pose.

The two structures revealed alternative noncovalent solutions for engaging the T790M/C797S ATP-binding pocket. HG-14-10-04 uses an extended and comparatively flexible architecture, whereas UCN-02 engages the pocket through a more rigid polycyclic scaffold and a distinct hydrophobic-contact network. These distinct binding modes provide complementary structural templates for structure-activity relationship studies. In particular, the solvent-exposed and weakly ordered terminal region of HG-14-10-04 may permit chemical modification, whereas the compact binding mode of UCN-02 identifies an alternative scaffold arrangement for optimizing pocket occupancy. Together, these structures provide a foundation for improving potency, mutant preference, and drug-like properties of these compounds for treating drug resistant NSCLC.

## Discussion

Mutant-selective compound binding is critical for improved efficacy and safety of small-molecule cancer therapies. Using triple-mutant EGFR as clinically relevant target, we demonstrated how prioritizing compounds solely by their predicted activity against mutant EGFR can overlook substantial WT EGFR inhibition, therefore providing an incomplete basis for selecting mutant-preferential candidates. To address this limitation, we developed a mutant-aware, paired activity machine-learning framework that considers mutant potency and WT activity together during early compound prioritization. Application to EGFR L858R/T790M/C797S recovered several compounds with reported activity against resistant EGFR variants^11,16,30,31^, supporting an overall agreement between the predictions and existing experimental evidence. We subsequently focused on compounds with remaining characterization gaps, particularly HG-14-10-04 and UCN-02.

Experimental profiling supported HG-14-10-04 as a mutant-preferential inhibitor and extended its previous biochemical characterization^16^ to mutant-WT comparisons, cellular activity and structural analysis. HG-14-10-04 showed consistent mutant preference both in biochemical inhibition (kinase assay) and apparent MST binding, and it inhibited cells expressing the triple-mutant EGFR, whereas WT-EGFR-amplified cells were less sensitive. UCN-02 retained a substantial WT activity and therefore a compromised selectivity profile. Our confirmatory assays were conducted at 1,000 µM ATP, well above the measured apparent ATP Kₘ of 6.7 µM. Assuming reversible ATP-competitive inhibition, the apparent IC_50_ values of 3.1 µM for HG-14-10-04 and 5.9 µM for UCN-02 correspond to Cheng–Prusoff-derived Kᵢ estimates of approximately 20.6 and 39.3 nM, respectively. These estimates are consistent with nanomolar inhibitory potency under the assumed mechanism, despite the micromolar IC₅₀ values measured at high ATP.

The co-crystal structures revealed two distinct noncovalent approaches to targeting the C797S mutated receptor. The HG-14-10-04 adopts an extended pose and forms two hinge hydrogen bonds with Met793, while a terminal substituent projects towards the solvent-exposed entrance of the ATP-binding pocket. The weaker electron density in this region is consistent with conformational flexibility and identifies a position that could be modified to improve solubility, cellular exposure or interactions with less-conserved residues without disrupting the hinge-binding core. The chlorine substituent of HG-14-10-04 is positioned near Met790, suggesting a possible contribution to recognition of the T790M-containing pocket, although this hypothesis requires direct testing. UCN-02 instead adopts a compact orientation imposed by its rigid polycyclic scaffold and occupies the hydrophobic region of the pocket through a broader contact network. These structures provide experimentally defined starting points for further optimization of potency and mutant preference. Further development would also require assessment of CNS exposure^33^, given the importance of intracranial activity in treating EGFR-mutant NSCLC.

In future, defining the target profile and patient population for mutation-specific therapies will become increasingly important, since C797S can occur in different mutational and allelic contexts, while resistance to EGFR-targeted therapies may arise through EGFR-independent mechanisms^34^. Although our future research will take paired-activity prediction frameworks further and accommodate more key variables, such as the CNS exposure, the next-generation EGFR inhibitors will compete not only with previous small-molecular inhibitors, but also with combination regimes, bispecific antibodies^35^, antibody-drug conjugates^36^ and therapies targeting bypass pathways such as MET^37^. Since these approaches may increasingly fragment the population available for mutation-specific therapies, future inhibitor development using AI/ML and paired-activity modelling frameworks will benefit from moving beyond optimization against a single resistance genotype towards compounds with broader coverage of relevant EGFR variants while passing CNS and sparing the wild-type receptor.

Although the present study is limited by the size and heterogeneity of the paired bioactivity dataset, and the prospective comparison between the individual modelling and prioritization components, our results demonstrate how paired mutant-WT modelling can incorporate selectivity into an early stage of compound prioritization. Expanding this approach to broader chemical libraries and additional EGFR genotypes and ultimately integrating properties such as CNS penetration and broader resistance coverage, can provide a possible route toward computational prioritization against the target profiles required for future generations of EGFR inhibitors. Finally, paired activity modelling provides an informed means to incorporate predicted mutant selectivity and differential activity into systematic candidate screening, and these findings support continued evaluation of the framework also for other cancer-driving, mutated disease targets beyond EGFR and drug-resistant NSCLC.

## Methods

### Study design

We developed a two-stage computational framework to identify compounds with preferential activity against mutant EGFR. The first stage used paired, continuous bioactivity measurements to estimate mutant activity, wild-type (WT) activity and mutant–WT activity difference. The second stage applied an independently trained classifier to provide complementary classification-based prioritization of compounds predicted to be specifically active against EGFR L858R/T790M/C797S. Model development and comparison were conducted using compound-level scaffold splits, so that compounds sharing a Bemis–Murcko scaffold were not distributed across both training and test sets^38,39^. Compounds satisfying all regression criteria and the mutant-classification criterion were subsequently consolidated by chemical identity and structural redundancy before proceeding to experimental evaluation.

### Curation of paired EGFR bioactivity measurements

Paired EGFR measurements were assembled from ChEMBL release 37^40^ and BindingDB^41^. A pair was defined as bioactivity measurements for the same standardized compound against WT EGFR and an EGFR variant, with compatible endpoint and assay annotations. Detailed distributions of data sources, activity endpoints, assay classes, confidence levels and variant types are provided in **Supplementary Table 2**.

Activities reported in molar units were converted to pActivity as

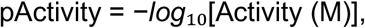

where Activity (M) is the reported IC_50_, K_d_ or K_i_ in molar units. Endpoint identity was retained as a categorical model input, rather than treating IC_50_, K_d_ and K_i_ as experimentally interchangeable quantities. For each pair, differential activity was represented as

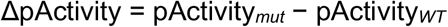

Thus, a positive value indicates preferential activity against the mutant; for measurements obtained under comparable conditions, the corresponding fold preference is 10^ΔpActivity. Duplicate records were reconciled using standardized chemical identity, EGFR variant, endpoint, assay metadata and source provenance. ΔpActivity was calculated for each paired measurement before aggregation. WT pActivity, mutant pActivity and paired ΔpActivity were then aggregated separately using their respective medians. Records with unresolved chemical structures, non-numeric activity values or incompatible paired annotations were excluded. All retained records preserved their source, assay type, endpoint, variant annotation and confidence metadata.

### Curation of the EGFR L858R/T790M/C797S classification dataset

The triple mutant-classification dataset comprised 2,099 compounds, including 997 experimentally measured active compounds, 339 experimentally measured inactive compounds, and 763 negative compounds randomly sampled from ChEMBL. Experimentally measured compounds with IC_50_ values below 50 nM were labelled as active, whereas those with IC_50_ values above 500 nM were labelled inactive. The 763 random compounds did not have experimentally measured activity against EGFR L858R/T790M/C797S and were included as presumed-negative examples for the classification workflow. The distribution of active and negative compounds across ChEMBL, BindingDB and the patent literature is reported in **Supplementary Table 3**.

### Triple mutant-classifier training and validation

Six classifier families were evaluated: support vector machine (SVM), logistic regression, random forest, extreme gradient boosting^42^ (XGBoost), k-nearest neighbours and a feed-forward artificial neural network. For each fingerprint–algorithm combination, the highest-ranked parameter setting from the cross-validation record was evaluated on common scaffold folds. Hyperparameters for each fingerprint–algorithm combination were selected by ten-fold cross-validation within the training set (**Supplementary Table 4**). The selected hyperparameter settings were then fixed and applied consistently across the same ten repeated scaffold-based five-fold partitions to ensure that all model comparisons used identical data splits. ROC area under the curve (ROC-AUC) and Matthews correlation coefficient (MCC) were the primary classification metrics; accuracy, balanced accuracy, precision and recall were secondary metrics. Reported confidence intervals and pairwise significance comparisons were calculated from matched seed-level means as described above.

### Compound standardization and molecular representations

Chemical structures were parsed and standardized with RDKit version 2023.09.5.^43^. Salts and disconnected components were resolved by retaining the largest organic fragment, charges were normalized where chemically appropriate, and structures were canonicalized. Connectivity-based identifiers were used for leakage auditing, whereas full InChIKeys were retained when stereochemical identity was required during candidate consolidation.

Seven two-dimensional compound representations were evaluated in the paired-regression benchmark: ECFP4 (Morgan radius 2, 2,048 bits), ECFP6 (Morgan radius 3, 2,048 bits)^21^, RDKit topological fingerprints (2,048 bits), hashed atom-pair fingerprints (2,048 bits), MACCS keys (167 bits), CDK Standard fingerprints (1,024 bits) and CDK ECFP6 fingerprints (1,024 bits)^22^. The precomputed CDK matrices were aligned to the same standardized compounds, endpoints and frozen folds used for the RDKit-derived representations. Fingerprint dimensionality and generation settings were held fixed across all model comparisons.

### Encoding of EGFR variants

EGFR variants were represented using three alternative encoding strategies: an explicit mutation descriptor, a feature-hashed mutation representation and a protein language-model representation. For the explicit mutation descriptor, each variant annotation was first decomposed into individual amino-acid substitutions, deletions and insertions. The resulting feature vector encoded the number and type of alterations, mutation positions, exon locations, and the identities of the WT and altered amino acids. This representation allowed the model to distinguish different EGFR variants directly, including single and compound mutations, without requiring the complete protein sequence as input.

For the feature-hashing representation, the complete standardized variant annotation was mapped to a 256-dimensional signed-hash vector^44^. Mutation count was included as an additional feature. This approach provided a fixed-length representation of each variant, but it did not explicitly preserve the biochemical meaning of individual amino-acid changes.

For the protein language-model representation, a mutant sequence was generated from the canonical human EGFR sequence (UniProt P00533)^45^ according to the standardized variant annotation. WT and mutant sequences were separately encoded using ESM-2 protein language-model, which generates a 640-dimensional embedding for each residue^20^. Two sequence summaries were calculated: a global embedding obtained by mean-pooling across the EGFR kinase domain (residues 650–1000), and a local embedding obtained by mean-pooling over the union of windows extending 16 residues on either side of each mutation site. Overlapping residues were counted only once.

To represent mutation-induced changes directly, the corresponding WT embedding was subtracted from each mutant embedding:

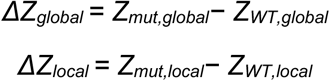

The global and local difference vectors were concatenated to form the ESM-difference representation. Within each cross-validation fold, principal-component analysis was fitted using the training partition only and reduced the concatenated representation to 16 components. The fitted transformation was subsequently applied without refitting to the corresponding validation and test partitions.

Variant annotations were classified into three reconstruction categories. Exact variants contained sufficient information to generate the mutant sequence unambiguously. Surrogate variants were represented using a standardized representative sequence when the original annotation described a recurrent alteration without specifying every residue; for example, a common exon 19 deletion was represented as E746_A750del. Unresolved variants could not be converted into a unique mutant sequence. These records were retained because they contained valid paired bioactivity measurements, but no sequence alteration was inferred. Their ESM-difference vectors were therefore set to zero, and a separate one-hot reconstruction-quality indicator identified them as unresolved. Thus, a zero ESM-difference vector for these records represented missing sequence-level information rather than biological equivalence between mutant and WT EGFR.

### Frozen scaffold-based cross-validation

Bemis–Murcko scaffolds were calculated from standardized structures^38^, producing 376 compound scaffold groups in the paired dataset^39^. Complete scaffold groups were assigned to five folds by sorting groups by size and greedily allocating each group to the least-populated fold; the random seed was used only to resolve ties. For each outer fold, one fold was held out for testing, the next fold cyclically was used for validation and the remaining three folds were used for training. No standardized compound or scaffold group was shared across the train-test folds. All representations and algorithms were evaluated on identical frozen assignments. Ten random seeds were evaluated: 104, 410, 853, 746, 732, 360, 156, 556, 450 and 727.

### Classical regression model baselines

Separate regressors were trained for predicting mutant pActivity, WT pActivity and direct *Δ*pActivity. The classical benchmark included the training-set mean predictor, ridge regression, elastic net, k-nearest-neighbor regression, support vector regression (SVR), random forest, extremely randomized trees and histogram gradient boosting. Specifically, ridge regression used α = 1; elastic net used α = 0.01, an L1 ratio of 0.5 and a maximum of 10,000 iterations; k-nearest neighbors used 10 distance-weighted neighbors; and RBF-SVR used C = 10, gamma=scale and ε = 0.1. Random forest and extremely randomized trees used 500 estimators, a minimum leaf size of 2 and square-root feature subsampling. Histogram gradient boosting used 300 boosting iterations, a learning rate of 0.05, 31 maximum leaf nodes, L2 regularization of 1 and early stopping.

The molecular-representation benchmark used a fixed regression algorithm and identical folds to isolate the effect of the fingerprints. The mutation-representation benchmark analogously fixed the compound representation and regression algorithm. Model performance was quantified by root-mean-square error (RMSE), mean absolute error, Pearson correlation and Spearman correlation between observed and out-of-fold predictions.

### Neural-network regression baselines

Two neural architectures were evaluated under the same frozen folds. The multitask multilayer perceptron received an ECFP6 vector, endpoint identity and a selected EGFR variant representation. The compound branch comprised layers of 512 and 256 units with layer normalization, GELU activation and dropout. The protein branch projected its input to 128 units. The WT head received the compound and endpoint representations, whereas the mutant and *Δ*pActivity heads additionally received the protein representation. The three heads were optimized jointly using mean-squared error. Targets were standardized using training-fold means and standard deviations and transformed back to the original scale for evaluation. Training used AdamW, a learning rate of 8 × 10^−4^, weight decay of 1 × 10^−4^, a batch size of 128 and early stopping with a patience of 30 epochs over a maximum of 250 epochs.

The graph model represented compounds as molecular graphs and used a four-layer graph isomorphism network with 192 hidden units, residual connections and layer normalization. Global mean pooling generated a 256-dimensional compound embedding, which was combined with the 128-dimensional protein branch and endpoint representation in the same task-specific output design. The graph model was trained with AdamW using a learning rate of 6 × 10^−4^, weight decay of 2 × 10^−4^, a batch size of 96, gradient clipping at 5 and the same early-stopping procedure.

### Statistical comparison of regression models

For each performance metric, values were first averaged across the five folds within a seed. The resulting ten seed-level estimates were treated as paired observations. Group summaries are reported as the mean and 95% confidence interval across seeds, calculated using the two-sided Student-t critical value. Pairwise comparisons shown in the figures were performed with two-sided non-parametric Wilcoxon signed-rank tests on matched seed-level values, followed by Holm correction for multiple comparisons within each comparison family. RMSE was the primary error measure and Pearson correlation was the primary linear association measure; the remaining metrics are reported as supporting analyses.

### Source-controlled validation of the mutant classifier

Because activity labels were unevenly distributed across data sources, two additional analyses were conducted. First, a source-balanced dataset was constructed independently for each seed by down-sampling the active and inactive classes within each source to the corresponding minority-class size; this produced balanced contributions of 137 active and 137 inactive ChEMBL compounds, 201 active and 201 inactive patent compounds, and 22 active and 22 inactive BindingDB compounds. The fixed ECFP6 RBF-SVM (C = 10, gamma=scale) was then evaluated by five-fold scaffold cross-validation. Second, 20 Y-randomization controls were generated by permuting activity labels within the source before applying the same source-balanced scaffold-validation procedure^46^. These analyses assessed whether predictive performance persisted after controlling source composition and whether it exceeded performance obtainable from randomized labels. Source-balanced and label-randomized performance estimates were compared using two-sided exact Mann–Whitney U tests, with Holm correction across ROC–AUC and MCC.

### Ensemble inference and compound-library screening

The virtual-screening input was a Taosu compound-structure archive^28^ (Shanghai Taosu Biochemical Technology Co., Ltd.). The source files contained 12,322,705 rows before filtering, and the compounds were standardized using the same chemical rules as the model-development datasets. For regression inference, ECFP6 was constructed from the RDKit Morgan radius-3, 2,048-bit representation. Mutant and WT IC₅₀ estimates were obtained from the corresponding predicted pActivity values. Selectivity was evaluated using both the direct ΔpActivity prediction and the difference between separately predicted mutant and WT pActivity. These were complementary estimates derived from the same training data, rather than statistically independent measurements. The classifier received its corresponding archived 1,024-bit ECFP6 representation. The decision-function score was used for classification and was not interpreted as a calibrated probability.

Compounds were retained only when all four regression-screening criteria were met: predicted mutant IC_50_ < 50 nM, predicted WT IC_50_ > 100 nM, direct predicted ΔpActivity > 0.699, corresponding to a predicted mutant preference of >5-fold and derived WT-to-mutant IC_50_ ratio >5. The WT threshold was used as a quantitative enrichment criterion and was not interpreted as the inactive-class boundary used by the separate classifier. Regression-selected compounds were then evaluated with the ECFP6-SVM mutant classifier and retained only when classified as active against EGFR L858R/T790M/C797S.

Exact standardized connectivity matches to either model-development dataset were removed before further prioritization. Prior reports were then reviewed to distinguish previously characterized inhibitors from compounds with remaining cellular or structural characterization gaps. TQB3804, OBX02-011, CZC-54252 and NVP-TAE684 were excluded because biochemical and cellular activity had already been reported in relevant C797S-containing or triple-mutant models. HG-14-10-04, has previously reported triple-mutant biochemical activity^16^, but was retained for further cellular and structural characterization. The remaining candidates were assessed for structural similarity, molecular connectivity, stereochemical redundancy and experimental tractability to select five representatives. Retention in this set did not imply absence of all prior biochemical evidence.

### Recombinant EGFR expression and purification

Recombinant kinase-domain constructs of WT EGFR and EGFR L858R/T790M/C797S (residues 696–1022) were expressed in Sf9 cells using a baculovirus system. Sf9 insect cells were infected at a density of 3 × 10⁶ cells/mL and cultured in Sf-900 III SFM medium (Gibco) at 27°C for 72 h. Cells were harvested by centrifugation (2,000 × g, 15 min) and resuspended in a lysis buffer (50 mM Tris-HCl, pH 8.0; 300 mM NaCl; 10% glycerol; 2 mM TCEP). Cells were disrupted by sonication, and the lysate was clarified by centrifugation at 40,000 × g for 45 min at 4°C.

Supernatants were loaded onto a Ni²⁺-NTA affinity column (Qiagen) equilibrated with lysis buffer supplemented with 10 mM imidazole. Bound proteins were washed extensively, then eluted with 300 mM imidazole in the buffer. Fractions containing EGFR kinase domain were pooled and further purified by size-exclusion chromatography (Superdex 200 Increase, GE Healthcare) in gel-filtration buffer (20 mM HEPES pH 8.0, 500 mM NaCl, 1 mM TCEP).

Protein purity was assessed by SDS–PAGE, and fractions with >95% purity were pooled and concentrated using Amicon Ultra centrifugal filters (Millipore). Protein concentration was determined by Bradford assay, and aliquots were flash-frozen in liquid nitrogen and stored at –80°C until use in enzymatic and binding assays.

### Enzymatic activity assays

The EGFR kinetic parameters reported were determined in triplicate using an ATP/NADH-coupled enzyme assay in a 96-well format, as previously described^8^. The reaction mixture contained 1.0 mg/mL BSA, 2 mM MnCl₂, 1 mM phosphoenolpyruvate (PEP; Sigma-Aldrich, catalog no. P7002), 2 mM TCEP, 100 mM MOPS (pH 7.5), 5 mM synthetic peptide substrate ENAEYLRVA, 0.5 mM NADH, and rabbit muscle pyruvate kinase/lactate dehydrogenase (Sigma-Aldrich, catalog no. P-0294) at 1/40 of the final reaction volume. To determine the Michaelis constant for ATP (Kₘ), reactions were initiated by adding ATP at varying concentrations. Steady-state initial velocities were calculated from the slopes of the absorbance traces at 340 nm and fitted to the Michaelis-Menten equation to determine Vmax and Kₘ. The resulting ATP-dependent reaction rates were fitted to the Michaelis–Menten equation, yielding an apparent ATP Km of 6.7 µM.

For the initial candidate evaluation, sunvozertinib, UCN-02, HG-14-10-04, GSK1904529A and Mps1-IN-2 were tested against EGFR^L858R/T790M/C797S^ over a concentration range of 0–64 µM at 500 µM ATP. EAI045 was included as a reference inhibitor. HG-14-10-04 and UCN-02 were subsequently evaluated in focused concentration–response experiments against both EGFR^L858R/T790M/C797S^ and wild-type EGFR. These confirmatory assays used an ATP concentration of 1,000 µM.

For each EGFR protein, an enzyme-containing control lacking peptide substrate was included to determine the background signal. The corresponding background value was subtracted from each measurement before normalization. Relative kinase activity was calculated by dividing the background-corrected signal at each compound concentration by the background-corrected signal of the matched DMSO-treated control. All conditions were measured in triplicate. Concentration–response curves were fitted using variable-slope nonlinear regression in GraphPad Prism, and apparent IC₅₀ values were obtained from the fitted curves.

Assuming ATP-competitive inhibition, apparent Ki values were estimated using the Cheng–Prusoff relationship:

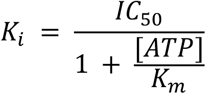

For the focused assays, an ATP concentration of 1,000 µM and the experimentally determined ATP Km of 6.7 µM were used. The apparent IC_50_ values of 3.1 µM for HG-14-10-04 and 5.9 µM for UCN-02 therefore corresponded to estimated apparent Ki values of approximately 20.6 nM and 39.3 nM, respectively.

### Cell culture and proliferation assays

#### Cell lines

Ba/F3 cells expressing EGFR L858R/T790M/C797S were maintained in RPMI-1640 with 10% FBS and 1 ng/mL IL-3. For experiments, IL-3 was withdrawn 24 h before drug treatment. H1975 (L858R/T790M), PC-9 (exon19 deletion), and A431 (wild-type amplification) were cultured in RPMI-1640 with 10% FBS, 2mM L-Glutamine and a mixture of penicillin-streptomycin. All cells were cultured at 37°C, 5% CO₂ and routinely tested for mycoplasma.

#### Drug treatment

HG-14-10-04, UCN-02, and EAI045 were dissolved in DMSO (10 mM stocks) and diluted to working concentrations (0.01–32 μM). Final DMSO did not exceed 0.1%. Cells were treated for 72 h.

#### Viability assay

H1975, PC-9 and A431 were seeded in 96-well plates at 3000 cell/well in 100ul complete culture medium. Cell viability was measured by MTT assay. Absorbance was measured at 570 nm with BioTek Cytation 5. Signals were normalized to DMSO control. Dose–response curves and IC_50_ values were generated in GraphPad Prism.

### Protein expression, crystallization, and structure determination

EGFR kinase-domain constructs comprising residues 696–1022 and harboring the T790M/C797S mutations were expressed in baculovirus-infected insect cells and purified as described previously^8^. To improve crystal quality and resolution, three substitutions were introduced into the activation loop (A-loop; E865A/E866A/K867A), as reported previously. Crystals were grown using the hanging-drop vapor-diffusion method. For co-crystallization, HG-14-10-04 was added to the purified protein at a final concentration of 0.5 ∼ 1 mM. The protein, at 3.5 mg/mL, was incubated with the inhibitor at 4 °C for 30 min before crystallization drops were set up. The reservoir solution contained 1.0 M potassium sodium tartrate, 0.2 M NaCl, 0.1 M HEPES (pH 8.5), and 5% glycerol. For UCN02 soaking experiments, the reservoir solutions used to grow inhibitor-free T790M/C797S crystals were 1.0 M potassium sodium tartrate, 0.2 M NaCl, 0.1 M HEPES (pH 8.5), and 5% glycerol. UCN02 was introduced by soaking the crystals for 3 h at room temperature in the corresponding crystallization buffer supplemented with 0.5∼1.0 mM. X-ray diffraction data were collected on beamlines BL18U1 and BL19U1 at the Shanghai Synchrotron Radiation Facility and processed using HKL3000. The structure was solved by molecular replacement in PHENIX, using the previously determined EGFR T790M/C797S_D3003 complex (PDB ID: 5ZTO) as the search model. CNS was subsequently used to perform simulated-annealing refinement and to calculate the electron-density maps, thereby reducing model bias and facilitating manual inspection and model adjustment. Several rounds of manual model building and crystallographic refinement were performed using COOT and PHENIX. The coordinates were modeled into the well-defined positive Fo-Fc electron density and included in subsequent refinement cycles. The corresponding topology and restraint parameter files were generated using PHENIX. Data-collection and refinement statistics are summarized in **Table 2**.

**Table 2.** Crystal structures of the EGFR T790M/C797S kinase domain bound to HG-14-10-04 or UCN-02.

| **Statistic** | **HG-14-10-04 complex** | **UCN-02 complex** |
| --- | --- | --- |
| PDB ID | 46VO | 46VR |
| **Data collection\*** |  |  |
| Space group | I23 | I23 |
| **Unit-cell dimensions** |  |  |
| a, b, c (Å) | 146.00, 146.00, 146.00 | 149.32, 149.32, 149.32 |
| $\alpha$, $\beta$, $\gamma$ (°) | 90.0, 90.0, 90.0 | 90.0, 90.0, 90.0 |
| Resolution (Å) | 103.23–2.89 (3.05–2.89) | 105.59–3.26 (3.66–3.26) |
| $R_{\text{pim}}$ | 0.093 (1.448) | 0.089 (1.393) |
| $I/\sigma(I)$ | 8.7 (1.4) | 12.5 (2.3) |
| Completeness (%) | 100 (100) | 100 (100) |
| Redundancy | 17.7 (18.1) | 34.3 (36.0) |
| **Refinement** |  |  |
| Resolution (Å) | 46.17–2.84 | 105.59–3.26 |
| No. of reflections | 11,657 | 7,909 |
| Rwork / Rfree | 0.2032 / 0.2488 | 0.2232 / 0.2746 |
| **No. of atoms** |  |  |
| Protein | 2,347 | 2,210 |
| Ligand/ion | 38 | 36 |
| Water | 0 | 0 |
| **Mean B factor (Å<sup>2</sup>)** |  |  |
| Protein | 77.28 | 89.92 |
| Ligand/ion | 85.20 | 102.14 |
| Water | 0 | 0 |
| **R.m.s. deviations** |  |  |
| Bond lengths (Å) | 0.010 | 0.008 |
| Bond angles (°) | 1.082 | 1.123 |
| **Ramachandran plot** |  |  |
| Favoured region (%) | 95.04 | 93.63 |
| Allowed region (%) | 4.61 | 6.37 |
| Outliers (%) | 0.35 | 0.0 |

\* Values in parentheses are for the highest-resolution shell.

## Code availability

The code used for data processing, model training, validation and computational screening, together with the fitted models and numerical results underlying the figures, is deposited at https://github.com/Jiezhu1995/EGFR-mutant-preference-ML.

## Acknowledgements

The authors thank Dr. Patrik Holm and Dr. Julia Lindqvist (Finnish Drug Discovery Center) for useful discussions and Drug Discovery and Chemical Biology Consortium (DDCB) core facility (HiLIFE-FIMM HTB unit), supported by the University of Helsinki and Biocenter Finland, for the compound testing. T.A. was supported by the Research Council of Finland (grants 373493 and 377685), the Cancer Society of Finland, the Norwegian Cancer Society (grants 216104 and 273810), Norwegian Health Authority South-East (grants 2020026 and 2023105), the Sigrid Jusélius Foundation, and iCAN – Digital Precision Cancer Medicine Flagship (iCAN-MULTIDRUG). J.Z. was supported by the China Scholarship Council (CSC), the Vilho, Yrjö and Kalle Väisälä Foundation, Biomedicum Helsinki Foundation and the Instrumentarium Science Foundation. H.Y. was supported by grants from K. Albin Johansson Stiftelse Foundation (No. 2023–12), the Maud Kuistila Memorial Foundation sr (No. 2024-0147B), the Finnish Cultural Foundation (No. 00251048), Biomedicum Helsinki foundation and the China Scholarship Council (No. 2020-I78). S.Z.J. was supported by the National Natural Science Foundation of China (No. 81903539) and the Innovation and Technology Program for Excellent Young Scholars of Higher Education of Shandong Province (No. 2022KJ145).

