## Supplementary Information for "Paired activity learning supports systematic identification of mutant-selective EGFR inhibitors"

**Supplementary Figures** Pages 2-9

**Supplementary Tables** Separate Excel file

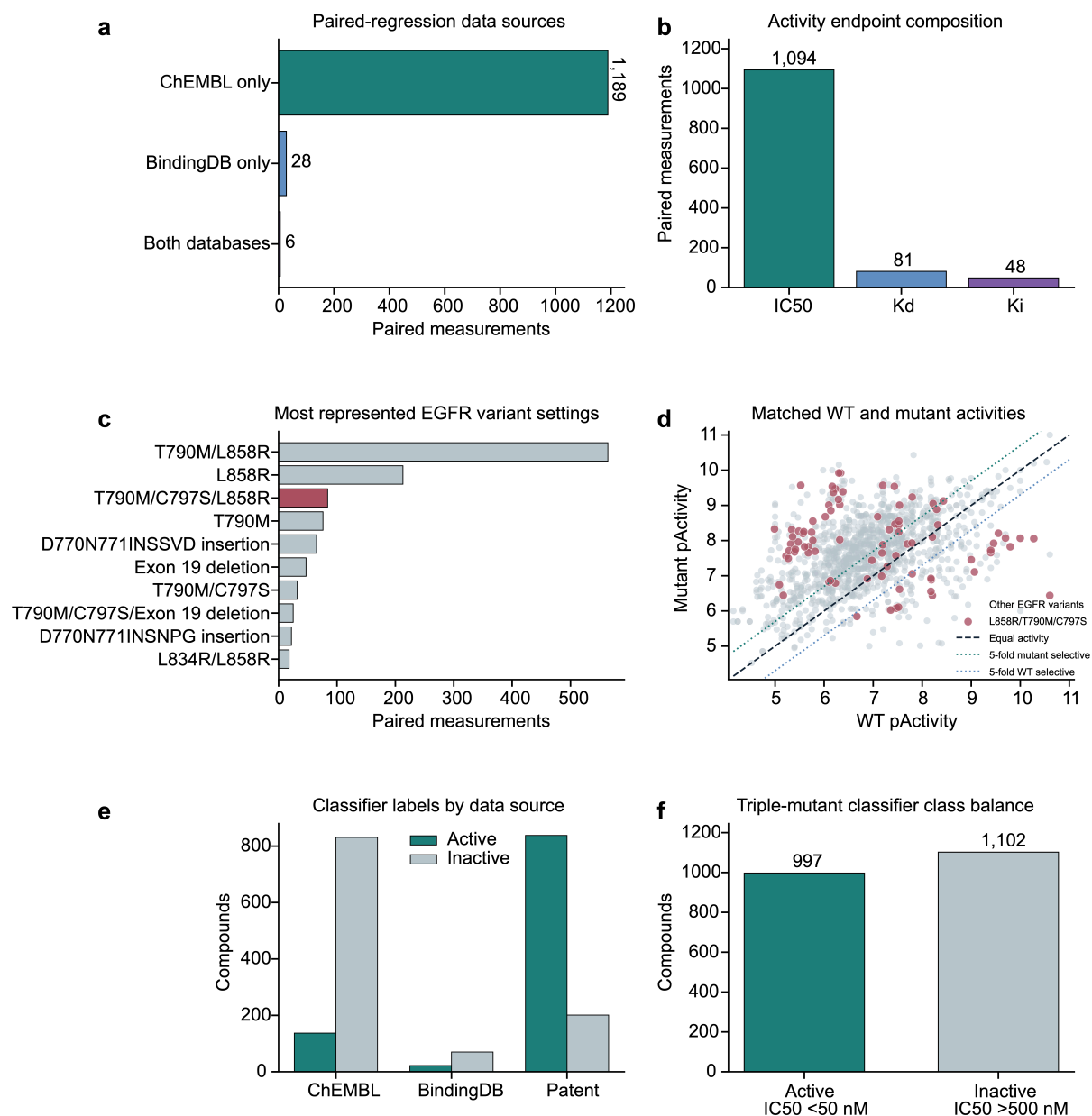

**Supplementary Figure 1 | Composition of the paired-regression and triple-mutant classification datasets.** **a**, Database provenance of the 1,223 paired WT-mutant activity measurements. Records occurring in both ChEMBL and BindingDB are shown as a separate, non-duplicated category. **b**, Distribution of activity endpoints in the paired-regression dataset: IC<sub>50</sub> (n = 1,094), K<sub>d</sub> (n = 81) and K<sub>i</sub> (n = 48). **c**, EGFR variant settings with the numbers of paired measurements. The paired dataset comprised 30 variant settings in total. **d**, Distribution of matched WT and mutant pActivity values. Colours distinguish EGFR L858R/T790M/C797S from other EGFR variants, and dashed lines indicate equal WT-mutant activity and 5-fold preference in either direction. **e**, Numbers of active and inactive triple-mutant classifier compounds obtained from ChEMBL, BindingDB and patent-derived records. **f**, Overall class balance of the EGFR L858R/T790M/C797S classifier dataset. The classifier dataset contained 997 experimentally measured active compounds, 339 experimentally measured inactive compounds and 763 presumed-negative compounds randomly sampled from ChEMBL. Measured active and inactive compounds were defined using IC<sub>50</sub> thresholds of <50 nM and >500 nM, respectively.

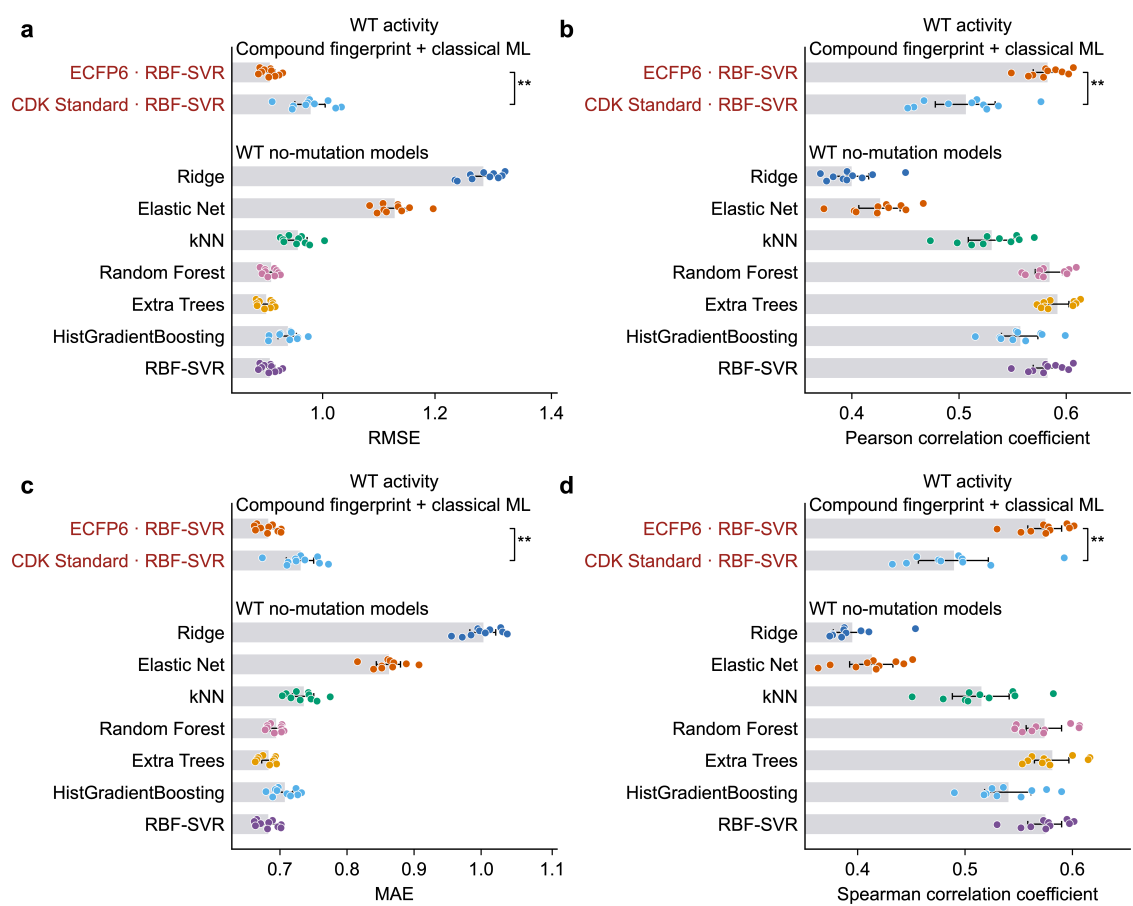

**Supplementary Figure 2 | WT EGFR activity prediction with ECFP6 fingerprints and without mutation encoding.** a-d, Performance of WT pActivity regression models evaluated using root mean square error (RMSE), Pearson correlation coefficient, mean absolute error (MAE) and Spearman correlation coefficient. The upper comparisons show ECFP6-RBF-SVR and CDK Standard-RBF-SVR. The lower comparisons show Ridge, Elastic Net, k-nearest neighbours (kNN), Random Forest, Extra Trees, HistGradientBoosting and RBF-SVR using ECFP6 without a mutation representation. Bars represent the mean of the seed-level five-fold averages across ten repeated scaffold partitions, error bars denote 95% confidence intervals and points denote individual seed-level averages. Red labels indicate the two highest-performing configurations for the corresponding endpoint and evaluation metric. Note that the scale of x-axis varies between the plots. ECFP6 and CDK Standard were compared using two-sided Wilcoxon signed-rank tests across matched seeds with Holm correction. \*$p < 0.05$, \*\*$p < 0.01$.

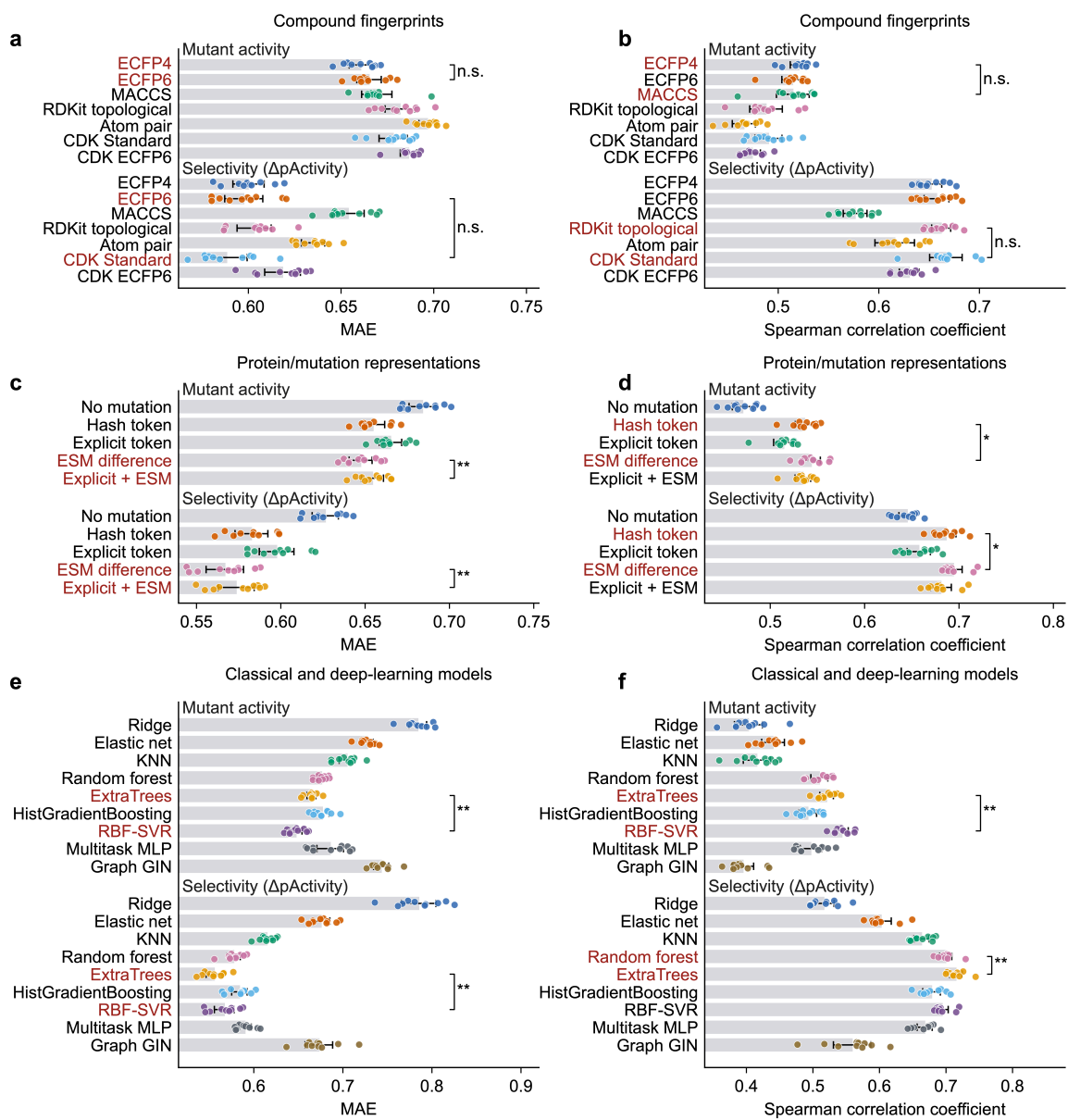

**Supplementary Figure 3 | Secondary metrics support the paired-regression representation benchmark.** **a,b**, Performance of seven compound fingerprints for mutant pActivity and ΔpActivity prediction using mean absolute error (MAE) and Spearman correlation coefficient. **c,d**, Performance of the no-mutation, feature-hashed mutation representation, explicit mutation descriptor, ESM-difference and explicit-plus-ESM EGFR variant representations, evaluated using MAE and Spearman correlation coefficient. **e,f**, Comparison of the seven conventional regression models, multitask MLP and graph-based GIN using MAE and Spearman correlation coefficient. Bars represent the mean of the seed-level five-fold averages across ten repeated scaffold partitions, error bars denote 95% confidence intervals and points denote individual seed-level averages. Red labels indicate the two highest-performing configurations for the corresponding endpoint and evaluation metric. Note that the scale of x-axis varies between the plots. Statistical comparisons used two-sided Wilcoxon signed-rank tests across matched seeds with Holm correction within each panel. \*$p < 0.05$, \*\*$p < 0.01$, \*\*\*$p < 0.001$; n.s., not significant.

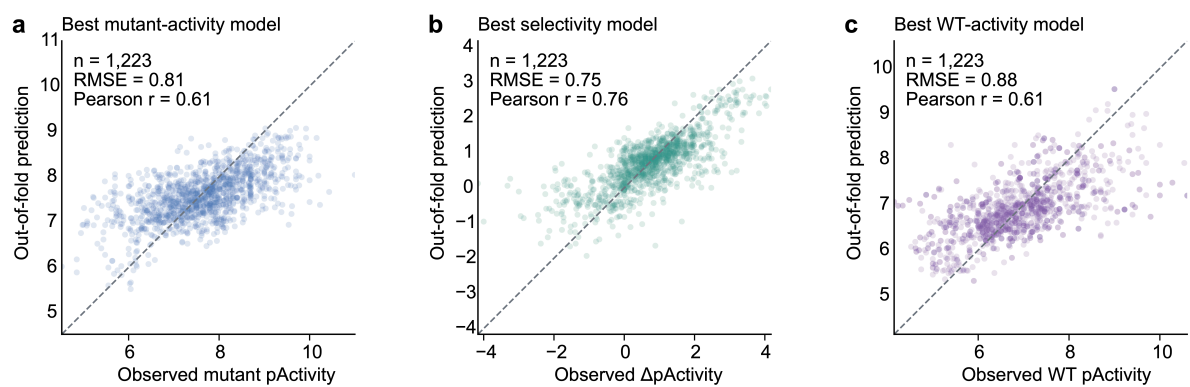

**Supplementary Figure 4 | Out-of-fold predictions of mutant activity, selectivity and WT activity.** **a**, Observed mutant pActivity versus out-of-fold predictions from the selected mutant-activity model (n = 1,223; RMSE = 0.81; Pearson r = 0.61). **b**, Observed ΔpActivity versus out-of-fold predictions from the selected selectivity model (n = 1,223; RMSE = 0.75; Pearson r = 0.76). **c**, Observed WT pActivity versus out-of-fold predictions from the selected WT-activity model (n = 1,223; RMSE = 0.88; Pearson r = 0.61). Dashed diagonal lines denote identity between observed and predicted values. Each point represents one paired WT–mutant measurement record. For each paired measurement record, the plotted prediction is the mean of the available out-of-fold predictions generated across the ten repeated scaffold partitions. The reported RMSE and Pearson correlation were recalculated from these averaged predictions and therefore differ from the mean seed-level metrics shown in Figure 2.

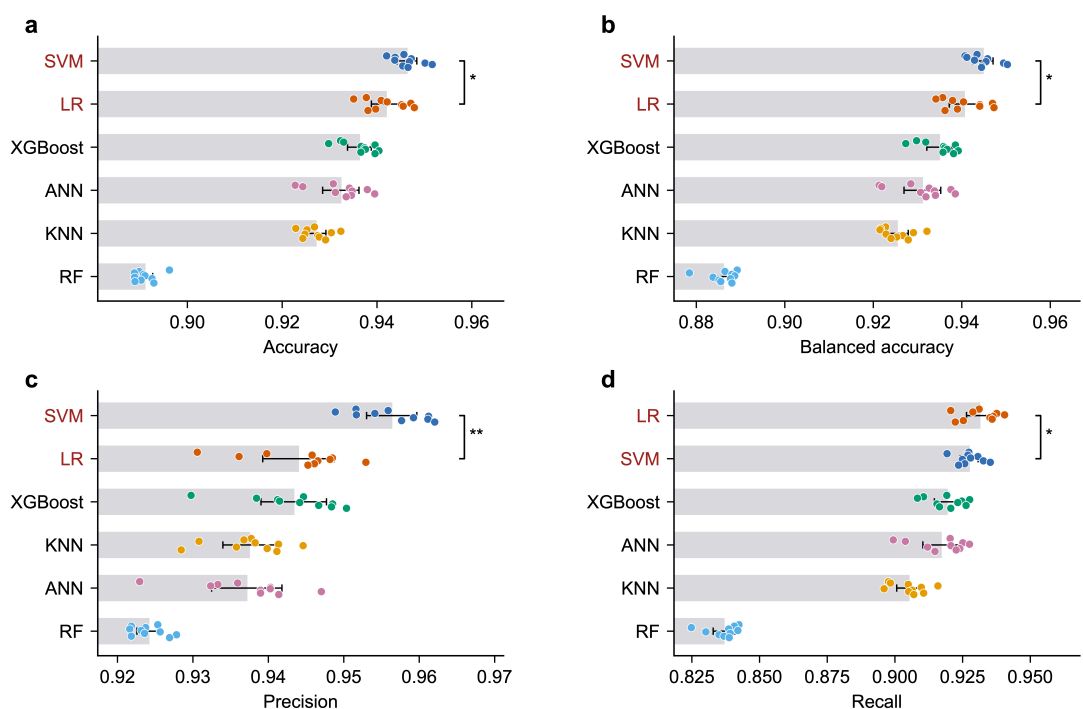

**Supplementary Figure 5 | Secondary classification metrics support selection of the ECFP6-SVM model.** a-d, Comparison of SVM, logistic regression (LR), XGBoost, artificial neural network (ANN), k-nearest neighbours (KNN) and random forest (RF) classifiers using accuracy, balanced accuracy, precision and recall. All models used ECFP6 fingerprints and were evaluated on the same ten repeated scaffold-based five-fold partitions. Each point represents the mean performance across the five test folds within one repeat. Bars show the mean of the ten repeat-level estimates, and error bars indicate 95% confidence intervals. The two highest-performing models in each panel were compared using two-sided Wilcoxon signed-rank tests on matched repeat-level estimates, followed by Holm correction across the four panels. \*$p < 0.05$, \*\*$p < 0.01$, \*\*\*$p < 0.001$.

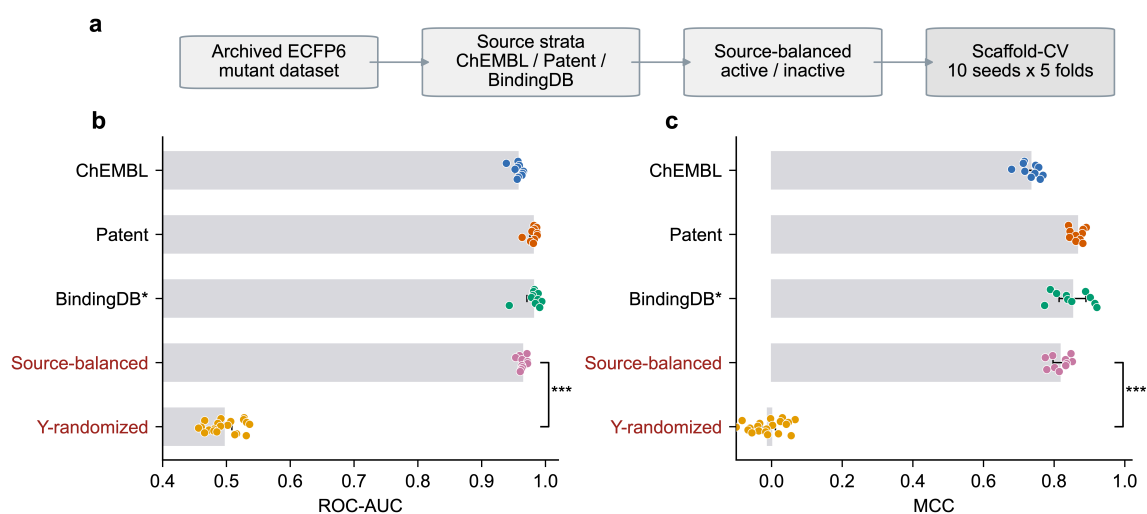

**Supplementary Figure 6 | Source-controlled scaffold validation supports activity-related learning beyond source imbalance.** **a**, Source-control workflow. The archived ECFP6 mutant dataset was stratified into ChEMBL, patent-derived and BindingDB subsets; active and inactive compounds were balanced within each source before repeated scaffold-based five-fold validation across ten seeds. **b,c**, Area under the receiving operating characteristic curve (ROC-AUC) and Matthews correlation coefficient (MCC), calculated from within-source scaffold validation, the combined source-balanced dataset and source-balanced Y-randomization. Bars represent means and 95% confidence intervals. Points represent ten seed-level five-fold means for observed activity labels or 20 independent label permutations for Y-randomization. The BindingDB subset contained 92 compounds and was considered exploratory. Source-balanced and Y-randomized estimates were compared using two-sided exact Mann–Whitney U tests, with Holm correction across ROC–AUC and MCC. \*\*\*$p < 0.001$.

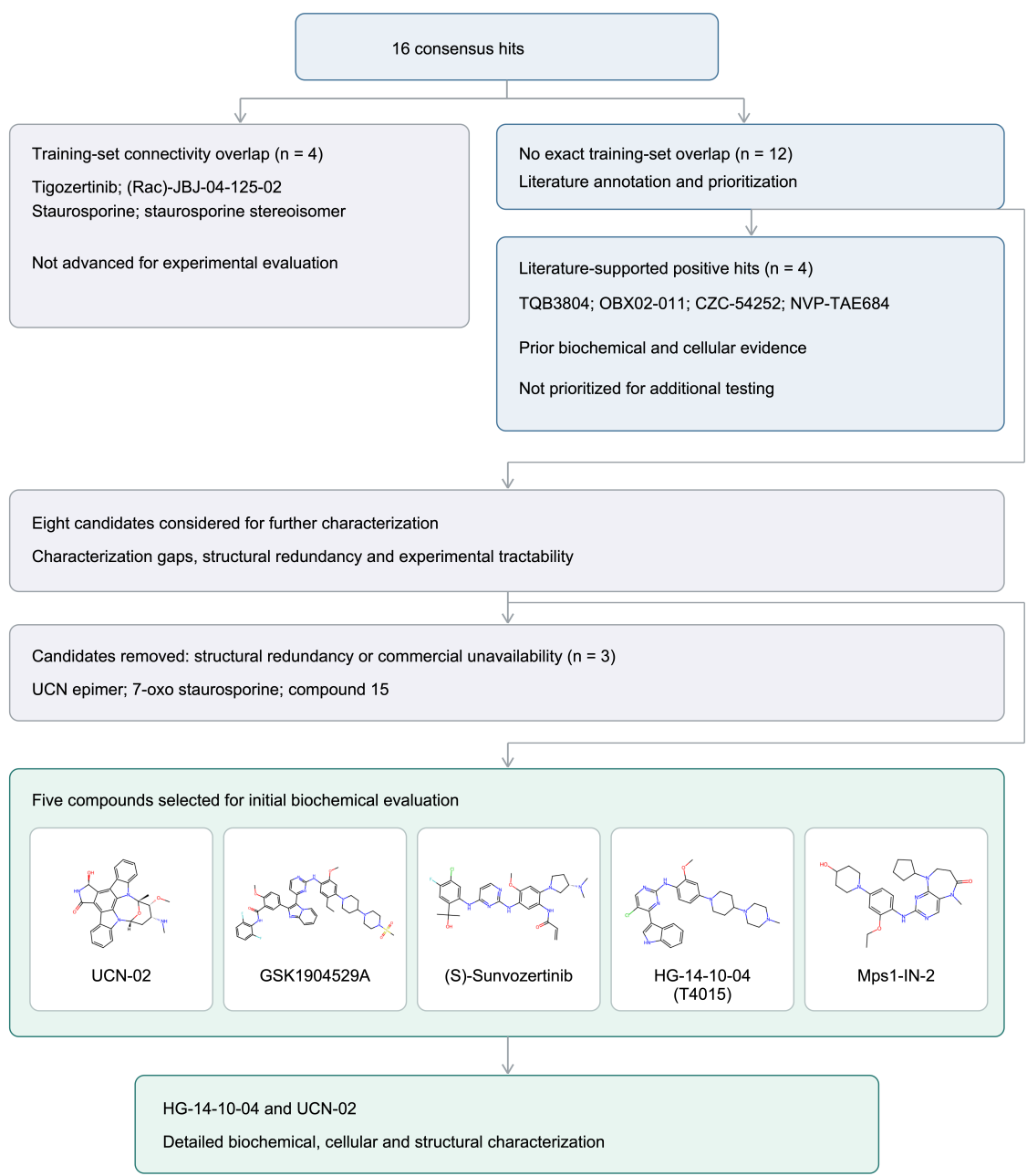

**Supplementary Figure 7 | Evidence annotation and experimental prioritization of consensus screening hits.** Of the 16 regression–classification consensus hits, four matched compounds in the model-development datasets and were not advanced for experimental evaluation. Among the remaining 12 compounds, four had reported biochemical and cellular activity in relevant C797S-containing or triple-mutant EGFR models and were retained as literature-supported positive hits but not prioritized for additional testing. Three further candidates were removed because of structural redundancy or commercial unavailability. The remaining five compounds underwent initial biochemical evaluation, followed by detailed biochemical, cellular and structural characterization of HG-14-10-04 and UCN-02. HG-14-10-04 had previously reported triple-mutant biochemical activity and was retained for further characterization rather than considered a newly discovered biochemical inhibitor.

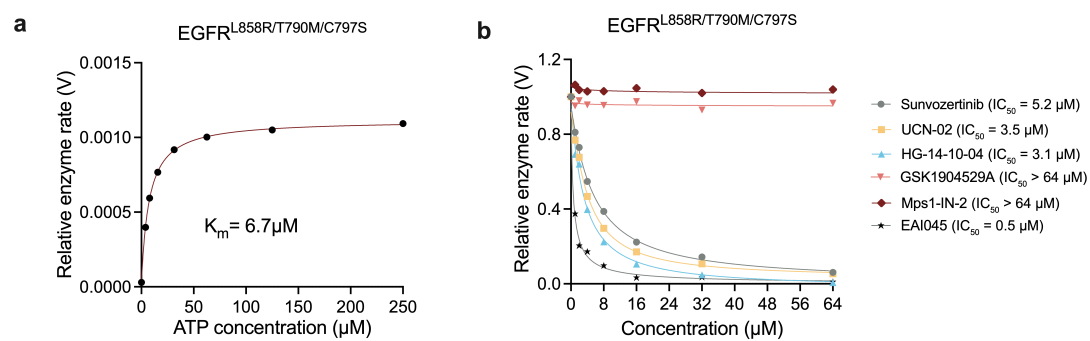

**Supplementary Figure 8 | Enzymatic characterization of candidate compounds.** **a**, ATP-dependent kinetics of recombinant EGFR L858R/T790M/C797S. Michaelis–Menten fitting yielded an apparent $K_m$ of 6.7 $\mu M$. **b**, Concentration-dependent effects of sunvozertinib, UCN-02, HG-14-10-04, GSK1904529A and Mps1-IN-2 on EGFR L858R/T790M/C797S enzyme activity. EAI045 was included as a reference inhibitor of the L858R/T790M mutant.
